# Analysis of SMAD1/5 target genes in a sea anemone reveals ZSWIM4-6 as a novel BMP signaling modulator

**DOI:** 10.1101/2022.06.03.494682

**Authors:** Paul Knabl, Alexandra Schauer, Autumn Penecilla Pomreinke, Bob Zimmermann, Katherine W. Rogers, Patrick Müller, Grigory Genikhovich

## Abstract

BMP signaling has a conserved function in patterning the dorsal-ventral body axis in Bilateria and the directive axis in anthozoan cnidarians. So far, cnidarian studies have focused on the role of different BMP signaling network components in regulating pSMAD1/5 gradient formation. Much less is known about the target genes downstream of BMP signaling. To address this, we generated a genome-wide list of direct pSMAD1/5 target genes in the anthozoan *Nematostella vectensis*, several of which were conserved in *Drosophila* and *Xenopus*. Our ChIP-Seq analysis revealed that many of the regulatory molecules with documented bilaterally symmetric expression in *Nematostella* are directly controlled by BMP signaling. Among the so far uncharacterized BMP-dependent transcription factors and signaling molecules we identified several, whose bilaterally symmetric expression may be indicative of their involvement in secondary axis patterning. One of these molecules, *zswim4-6*, encodes a novel nuclear modulator of the pSMAD1/5 gradient potentially promoting BMP-dependent gene repression. Strikingly, overexpression of the zebrafish homologue *zswim5* suggests that its effect on the pSMAD1/5 gradient is conserved between anthozoan Cnidaria and Bilateria.

## Introduction

The clade Bilateria unites animals with bilaterally symmetric body plans that are determined by two orthogonally oriented body axes, termed the anterior-posterior (A-P) axis and the dorsal-ventral (D-V) axis. These body axes form a Cartesian coordinate system, in which the location of different morphological structures is specified by gradients of morphogen signaling (Niehrs, 2010). Notably, bilaterality is also observed in representatives of a single animal clade outside of Bilateria: their evolutionary sister group Cnidaria (Berking, 2007; Finnerty et al., 2004). Common to all cnidarians is the formation of an oral-aboral (O-A) body axis which is patterned by Wnt/β-catenin signaling (Kraus et al., 2016; Lebedeva et al., 2021; Lee et al., 2007; Marlow et al., 2013; Momose et al., 2008; Momose and Houliston, 2007; Wikramanayake et al., 2003). However, in contrast to Medusozoa (jellyfish and hydroids), Anthozoa (sea anemones and corals) have an additional secondary, “directive” body axis (Fig. 1A), which is patterned by bone morphogenetic protein (BMP, Fig. 1B) signaling (Finnerty et al., 2004; Saina et al., 2009). BMP signaling has also been shown to regulate patterning of the D-V axis in Bilateria (Arendt and Nubler-Jung, 1997; Holley et al., 1995; Kozmikova et al., 2013; Lapraz et al., 2009; Özuak et al., 2014; van der Zee et al., 2006), however, it remains unclear whether this second, BMP-dependent body axis was a feature of the last common cnidarian-bilaterian ancestor and lost in Medusozoa, or whether it evolved independently in anthozoan Cnidaria and in Bilateria (Genikhovich and Technau, 2017). To address this fundamental evolutionary question, we need to gain a better understanding of how the directive axis is established and patterned.

**Figure 1.**
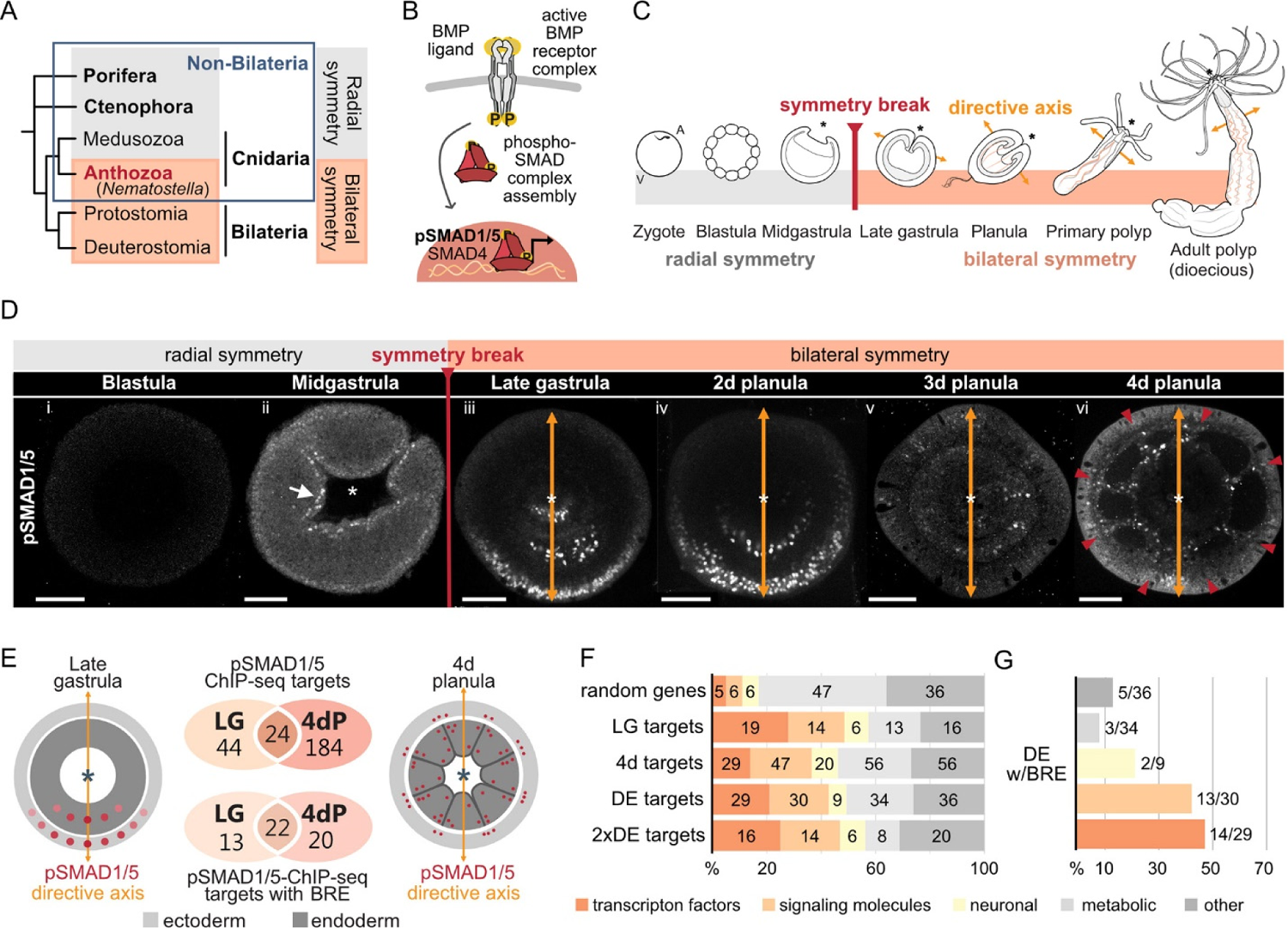
Bilateral body symmetry of the non-bilaterian sea anemone *Nematostella* is BMP signaling-dependent. (A) Bilateral body symmetry is observed in Bilateria and in anthozoan Cnidaria. (B) BMP signaling is initiated by BMP ligands binding to BMP receptors that trigger phosphorylation, assembly, and nuclear translocation of a pSMAD1/5/SMAD4 complex to regulate gene expression. (C) A BMP signaling-dependent symmetry break at late gastrula stage results in the formation of the secondary (directive) body axis in the sea anemone *Nematostella*. (D) BMP signaling dynamics during *Nematostella* development. No pSMAD1/5 is detectable in the blastula (Di). Nuclear pSMAD1/5 is localized in the blastopore lip of midgastrula (Dii), forms a gradient along the directive axis in the late gastrula (Diii) and 2d planula (Div). By day 3, the gradient progressively disperses (Dv), and the signaling activity shifts to the eight forming endodermal mesenteries (Dvi) and to the ectodermal stripes vis-à-vis the mesenteries (arrowheads). Images Dii-Dvi show oral views (asterisks). Scale bars 50 µm. (E) Comparison of the direct BMP signaling targets at late gastrula (LG) and 4d planula (4dP) shows little overlap. Schemes show oral views of a late gastrula and a 4d planula with red spots indicating the position of pSMAD1/5-positive nuclei in the ectoderm (light-grey) and endoderm (dark-grey). (F) Transcription factors, signaling molecules, and neuronal genes are overrepresented among the pSMAD1/5 targets compared to the functional distribution of 100 random genes. LG – late gastrula targets, 4dP – 4d planula targets, DE – pSMAD1/5 ChIP targets differentially expressed in BMP2/4 and/or GDF5-like morphants (padj ≤ 0.05), 2xDE targets - pSMAD1/5 ChIP targets differentially expressed in BMP2/4 and/or GDF5-like morphants (padj ≤ 0.05) showing ≥2-fold change in expression. (G) Fractions of each functional category of the differentially expressed pSMAD1/5 target genes (see panel F) containing BRE.

In contrast to Bilateria, where the D-V axis and the A-P axis usually form simultaneously and very early during development, the directive body axis of the sea anemone *Nematostella vectensis* appears only at gastrula stage (Matus et al., 2006a; Matus et al., 2006b; Rentzsch et al., 2006), while the O-A axis is maternally determined (Lee et al., 2007). The expression of the core components of the BMP signaling network *bmp2/4* and *chordin* is initially controlled by β-catenin signaling and radially symmetric (Kirillova et al., 2018; Kraus et al., 2016; Rentzsch et al., 2006). During gastrulation, the embryo undergoes a BMP signaling-dependent symmetry break establishing the directive axis at a molecular level (Rentzsch et al., 2006; Saina et al., 2009), with a BMP signaling activity gradient forming as revealed by antibody staining against phosphorylated SMAD1/5 (pSMAD1/5) (Fig. 1C-D) (Genikhovich et al., 2015; Leclère and Rentzsch, 2014). Each end of the directive axis expresses a set of BMP ligands and BMP antagonists: *bmp2*/*4* and *bmp5-8* are transcribed on the low-BMP signaling activity side of the directive axis together with the antagonist *chordin*, while the BMP ligand *gdf5-like* (*gdf5-l*) and the BMP antagonist *gremlin* are expressed on the high-BMP signaling activity side (Genikhovich et al., 2015; Rentzsch et al., 2006). The pSMAD1/5 gradient is maintained by the genetic interactions between these molecules, with Chordin likely acting as a shuttle for BMP2/4/BMP5-8 and Gremlin serving as a primary GDF5-like antagonist (Genikhovich et al., 2015). Another essential player in this network is the *repulsive guidance molecule (rgm)*, which is necessary for the maintenance of the low-BMP signaling side of the directive axis (Leclère and Rentzsch, 2014).

The graded BMP signaling activity is essential for the patterning of the endoderm and the formation of the so-called mesenteries, gastrodermal folds compartmentalizing the endoderm of the late planula into eight distinct chambers as demonstrated by knockdown (KD) experiments (Genikhovich et al., 2015; Leclère and Rentzsch, 2014). More specifically, complete abolishment of the BMP signaling gradient by KD of *bmp2/4*, *bmp5-8* or *chordin* results in embryos which are not only molecularly, but also morphologically radialized, failing to form any mesenteries (Genikhovich et al., 2015; Leclère and Rentzsch, 2014). In line with this, KD of *gdf5-l* or *gremlin* alter the profile of the BMP signaling gradient, and also lead to abnormal mesentery formation (Genikhovich et al., 2015).

Despite these important insights over the last decades, our knowledge of BMP-dependent directive axis patterning mechanisms, in particular regarding effector molecules linking BMP signaling and subsequent morphological bilaterality in *Nematostella* remains incomplete, precluding proper comparison of anthozoan directive and bilaterian D-V axis patterning. To address this, we performed a genome-wide search for direct BMP signaling targets at two developmental stages in *Nematostella* using ChIP-Seq with an αpSMAD1/5 antibody. We demonstrate that regulatory genes, including many with previously documented bilaterally symmetric expression, are overrepresented among direct BMP signaling targets. We also identify multiple previously uncharacterized transcription factors and signaling molecules, whose bilaterally symmetric expression suggests that these direct BMP signaling targets may be involved in the patterning of the directive axis and in endoderm compartmentalization. Several of these seem to be shared between *Nematostella*, *Drosophila*, and *Xenopus* as shown by comparison of pSMAD1/5 ChIP-Seq targets at similar developmental stages. Among the targets with maximum ChIP enrichment, we find *zswim4-6*, a gene encoding a so far uncharacterized zinc-finger protein with a SWIM domain (ZSWIM4-6), whose paralogs are also pSMAD1/5 targets in the frog. Functional analyses show that *Nematostella* ZSWIM4-6 can modulate the shape of the pSMAD1/5 gradient and appears to promote BMP signaling-mediated gene repression. Overexpression of the zebrafish paralogue *zswim5* revealed similar effects on the BMP signaling gradient in *Danio rerio*, indicating that the function of ZSWIM4-6 may be conserved in Bilateria.

## Results

### Identification of BMP signaling targets in *Nematostella*

Staining with an antibody recognizing the transcriptionally active phosphorylated form of SMAD1/5 (pSMAD1/5) demonstrates that BMP signaling activity is highly dynamic during early development in *Nematostella*. In the early *Nematostella* gastrula, nuclear pSMAD1/5 can be detected in a radially symmetric domain around the blastopore (Fig. 1D panel ii). Subsequently, it becomes restricted to one side of the gastrula (Fig. 1D panels iii-iv) in a symmetry breaking process, which has been shown to depend on BMP signaling itself (Saina et al., 2009). The resulting BMP gradient is crucial for the formation of the directive axis. The gradient, however, disappears by late planula, when the general patterning of the directive axis is complete, and the BMP signaling activity is confined to the mesenteries, and to eight stripes following the mesenteries in the ectoderm and merging in pairs to surround the future tentacle buds and ending in a circumoral ring (Fig. 1D panels v-vi, Supplementary video 1).

Although the role of BMP signaling in symmetry breaking and compartmentalization of the *Nematostella* endoderm has been well described, insight regarding direct targets of BMP signaling during these fundamental developmental processes is currently lacking. To generate a genome-wide list of direct BMP signaling targets in *Nematostella*, we performed ChIP-Seq using an anti-pSMAD1/5 antibody at two developmental stages: late gastrula, when BMP signaling forms a gradient along the directive axis (Fig. 1D panel iii), and late planula (4 d) when the mesenteries have formed and BMP signaling is confined to the mesenteries and to stripes of cells in the body wall ectoderm following the mesenteries (Fig. 1D panel vi). We found 68 direct pSMAD1/5 target genes at late gastrula stage and 208 in 4d planulae, 24 of which were bound by pSMAD1/5 at both developmental stages (Fig. 1E). MEME-ChIP analysis (Ma et al., 2014) of the sequences occupied by pSMAD1/5 showed that the BMP response element (BRE) sequence, a vertebrate SMAD1/5 binding motif, was enriched at many of the bound sites (Fig. 1E). Most genes identified as pSMAD1/5 targets at both stages carry a BRE motif (22/24), with 13 additional gastrula-specific targets and 20 additional 4d planula-specific targets containing a BRE site. Next, we manually annotated pSMAD1/5 targets using reciprocal best BLAST hits and found among them many key regulatory genes whose dependence on BMP signaling has previously been documented in *Nematostella* (Genikhovich et al., 2015; Leclère and Rentzsch, 2014; Saina et al., 2009). These included *gbx*, *hoxB*, *hoxD* and *hoxE* - i.e. genes shown to play a central role in the subdivision of the directive axis into mesenterial chambers and specifying the fate of each mesenterial chamber (He et al., 2018) - as well as the BMP signaling regulators *chordin*, *gremlin* and *rgm* (Genikhovich et al., 2015; Leclère and Rentzsch, 2014; Saina et al., 2009). Intriguingly, none of the BMP ligand-coding genes, i.e. *bmp2/4*, *bmp5-8* or *gdf5-like*, was found as a direct BMP signaling target (Supplementary Data 1).

To expand this analysis and gain insight into the response of the 252 newly found putative direct pSMAD1/5 target genes to abolished or reduced BMP signaling, we injected previously characterized antisense morpholino oligonucleotides against *bmp2/4* and *gdf5-like* (Genikhovich et al., 2015; Saina et al., 2009), as well as a standard control morpholino, and analyzed by RNA-Seq whether the pSMAD1/5 target genes were differentially expressed upon *gdf5-like* or *bmp2/4* KD (Supplementary Fig. 1, Supplementary Data 1). 139 direct pSMAD1/5 target genes were among the genes differentially expressed in the KD (p_adj_ ≤ 0.05), 64 of which showed more than a two-fold change in expression upon BMP2/4 and/or GDF5-like KD (Supplementary Data 1). The transcriptional response of the previously described downstream targets of BMP signaling confirmed previous *in situ* hybridization analysis, i.e. *chordin* and *rgm* were upregulated while *gbx*, *hox* genes and *gremlin* were downregulated in our RNA-Seq datasets confirming their specificity. Notably, out of 55 pSMAD1/5 target genes with BRE motifs, 37 (67.3%) were shown to be differentially expressed upon BMP2/4 and/or GDF5-like KD.

To gain insight into the function of pSMAD1/5 target genes in *Nematostella*, we roughly categorized them according to the putative function of their bilaterian homologs. We found that the fraction of transcription factors and signaling molecules was strongly increased among the direct BMP signaling targets at both developmental stages in comparison to 100 randomly selected genes from the *Nematostella* genome (Fig. 1F-G). This suggests that direct downstream targets of BMP signaling constitute a second tier of a regulatory cascade governing patterning and morphogenesis of the *Nematostella* gastrula and planula. Since we were particularly interested in identifying new players in the BMP-dependent regulation of the directive axis, we characterized the expression domain of a subset of pSMAD1/5 targets with so far unknown expression patterns and validated the expression of several previously described pSMAD1/5 targets using whole mount *in situ* hybridization (WISH). In accordance with a potential role in the directive axis regulation, we found several of the newly identified pSMAD1/5 targets to be bilaterally expressed (e.g. *otxB*, *tbx2/3*, *tbx20.1*, *p63*, *dusp1*, *bmprII*, *c-ski*, *morn*, *pik, zswim4-6*) (Fig. 2), further motivating the investigation of their function in directive axis patterning.

**Figure 2.**
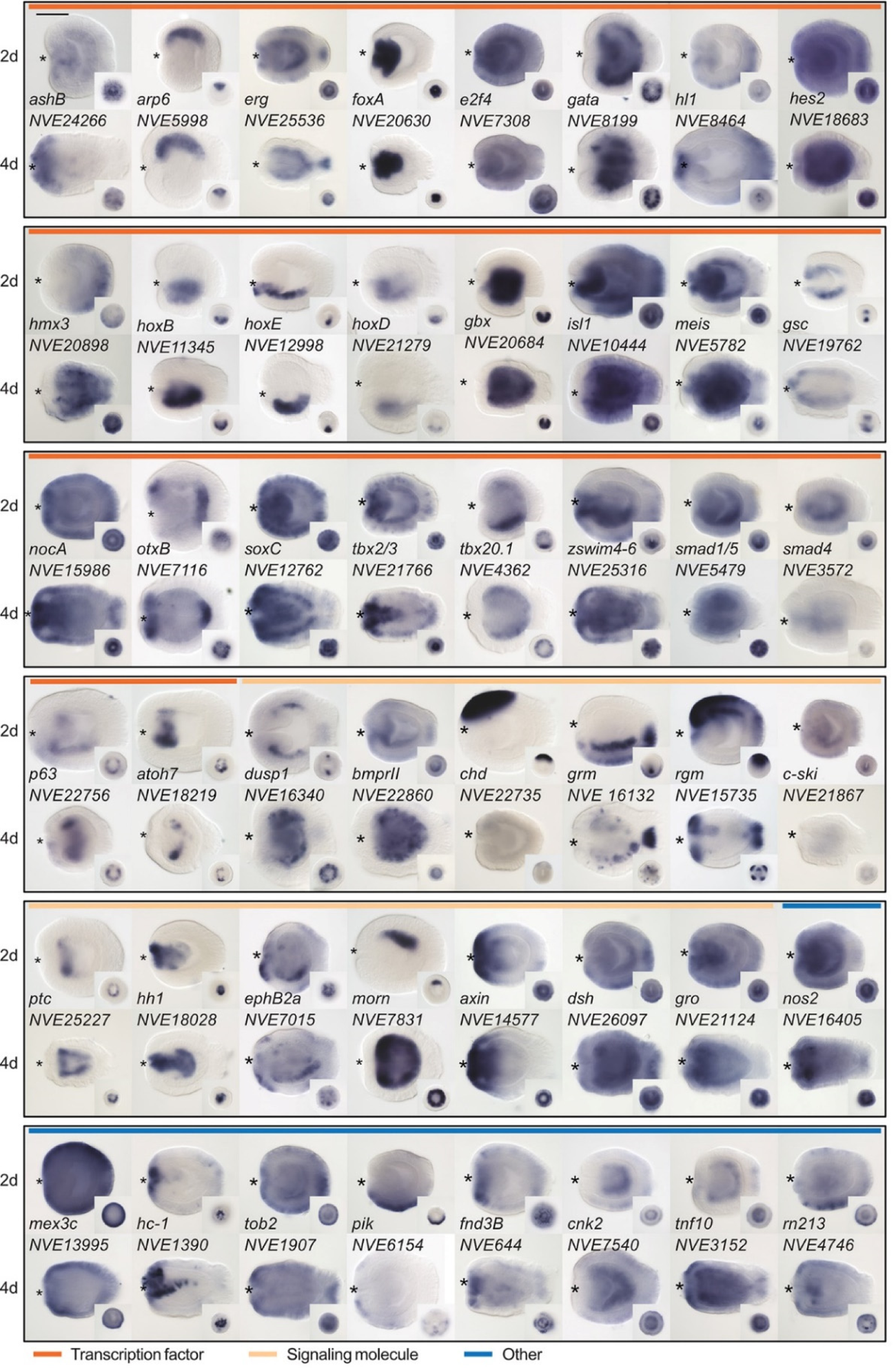
Expression patterns of a selection of the direct targets of BMP signaling in 2d and 4d planulae. In lateral views, the oral end is marked with an asterisk, inlets show oral views. Scale bar represents 100 µm.

### Conservation of direct BMP signaling targets between Anthozoa and Bilateria

To identify BMP signaling targets conserved between Anthozoa and Bilateria, we compared *Nematostella* pSMAD1/5 targets with available ChIP-Seq data from two bilaterians. In *Drosophila melanogaster*, pMAD targets were identified at two developmental time points (2 and 3 h after fertilization) (Deignan et al., 2016), and in *Xenopus laevis*, SMAD1 targets were found post-gastrulation (NF20 stage) (Stevens et al., 2017). In a 3-way comparison of *Nematostella* (*Nve*), fly (*Dme*), and frog (*Xla*), putative orthologs were identified using best reciprocal BLAST hits based on the bit score. We found 103 direct BMP signaling targets that were conserved between at least two organisms, of which four were shared by all three species (Fig. 3E). Anchoring the analysis on *Nematostella* indicates that several of the targets, which are shared between only two species in a strict 3-way comparison, can be assigned to all three species thus increasing the number of common targets to nine (*meis, gata, hoxB, tbx2/3, nkain, irx, zfp36l1, ptc, tp53bp2)*. Among the shared targets, conserved transcription factors (TFs) and signaling molecules (SMs) are enriched compared to genes with other functions (69/103). Multiple Hox genes are direct targets of BMP signaling in embryos of *Nematostella* (*hoxB, hoxD, hoxE*), fly (*dfd, pb, antp*) and frog (*hox3-7/9-11/13, hoxa1-2, hoxb1-9, hoxc3-6/8-12, hoxd1/3/4/8-11/13*); however, the orthology of the cnidarian and bilaterian Hox genes is currently unclear, due to the likely independent diversification of the “anterior” and “non-anterior” Hox genes in these two sister clades (Chourrout et al., 2006; Genikhovich and Technau, 2017).

**Figure 3.**
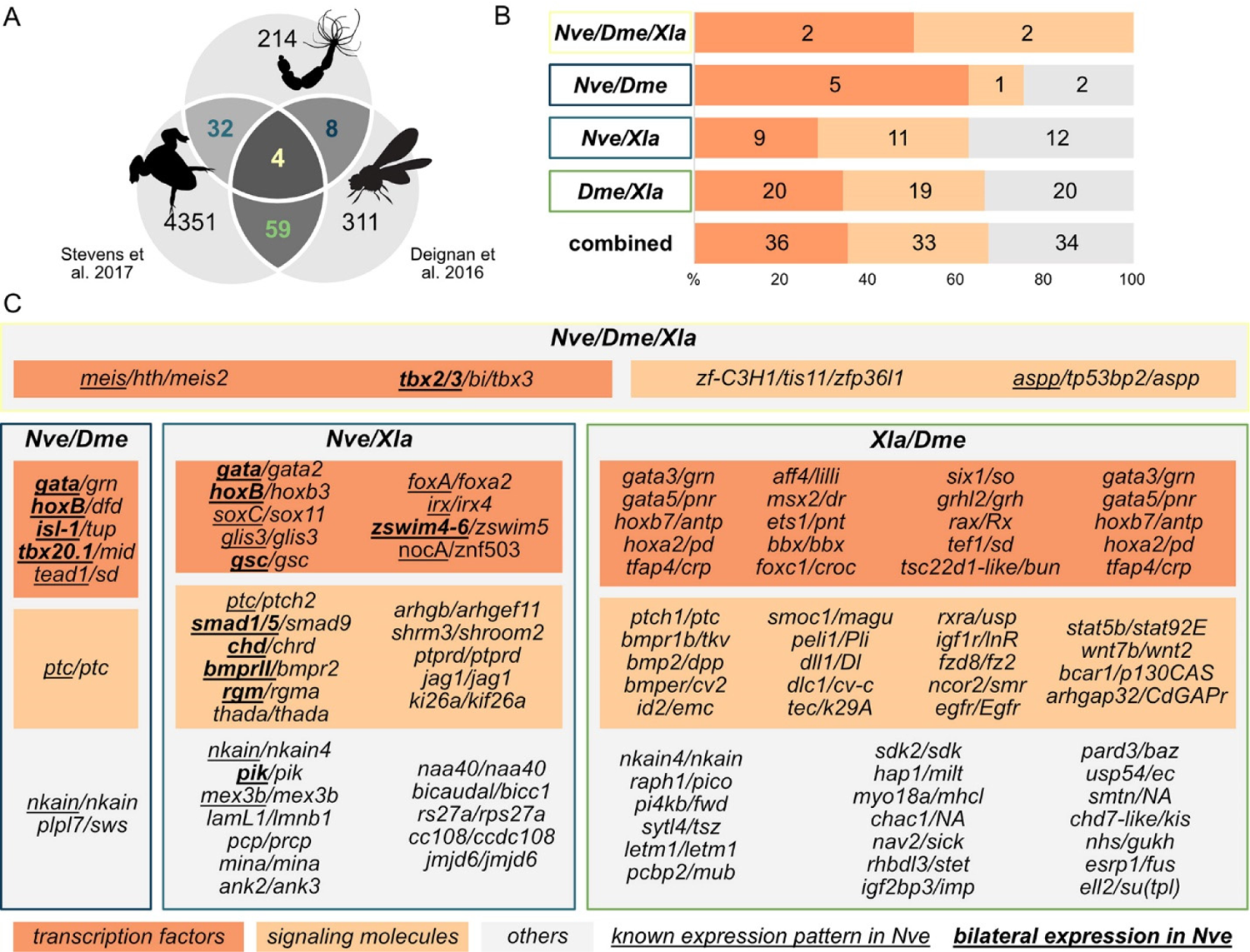
*Nematostella* and Bilateria share BMP signaling targets, which predominantly encode transcription factors and signaling molecules. (A) Overlap of BMP signaling targets at comparable embryonic stages in the 3-way comparison of *Nematostella*, *Drosophila* (Deignan et al., 2016) and *Xenopus* (Stevens et al., 2017). Orthology links were deduced by NCBI BLASTP of the respective proteomes with a cut-off e-value ≤1e-5, and reciprocal best BLAST hits were determined using the bit score. (B) Transcription factors and signaling molecules represent more than 60% of pSMAD1/5 targets shared between *Nematostella* (*Nve*), fly (*Dme*), and frog (*Xla*). (C) Gene names of orthologous targets shared between *Nematostella*, fly and frog. For targets shared with *Nematostella*, genes with known expression patterns in the embryo are underlined, genes expressed asymmetrically along the directive axis are underlined and bold.

### *zswim4-6* – a previously uncharacterized, asymmetrically expressed gene encoding a nuclear protein

One of the highest enrichment levels in our ChIP-Seq was detected for *zswim4-6* – a pSMAD1/5 target shared between *Nematostella* and *Xenopus* (Fig. 3). This gene encodes a zinc-finger containing protein with a SWIM-domain clustering together with ZSWIM4/5/6 of zebrafish (*Danio rerio*) and mouse (*Mus musculus*) (Fig. 4A). *Zswim4-6* starts to be expressed at the early gastrula stage around the blastopore (Fig. 4B-C’), concomitant with the onset of BMP signaling activity. At late gastrula, when a symmetry break in the BMP signaling activity establishes the directive axis, *zswim4-6* expression becomes restricted to the ectoderm and endoderm on one side of it (Fig. 4D-D’). Double *in situ* hybridization of *chd* and *zswim4-6* shows that *zswim4-6* is expressed opposite to *chd*, i.e. on the side of high pSMAD1/5 activity (Fig. 4E-E’), suggesting that BMP signaling activates *zswim4-6* expression. At late planula stage, *zswim4-6* expression is confined to the eight mesenteries (Fig. 4F-F’). Taken together, *in situ* hybridization analysis showed that the *zswim4-6* expression domain followed the dynamic changes in pSMAD1/5 activity (compare Fig. 1D and Fig. 4B-F’).

**Figure 4.**
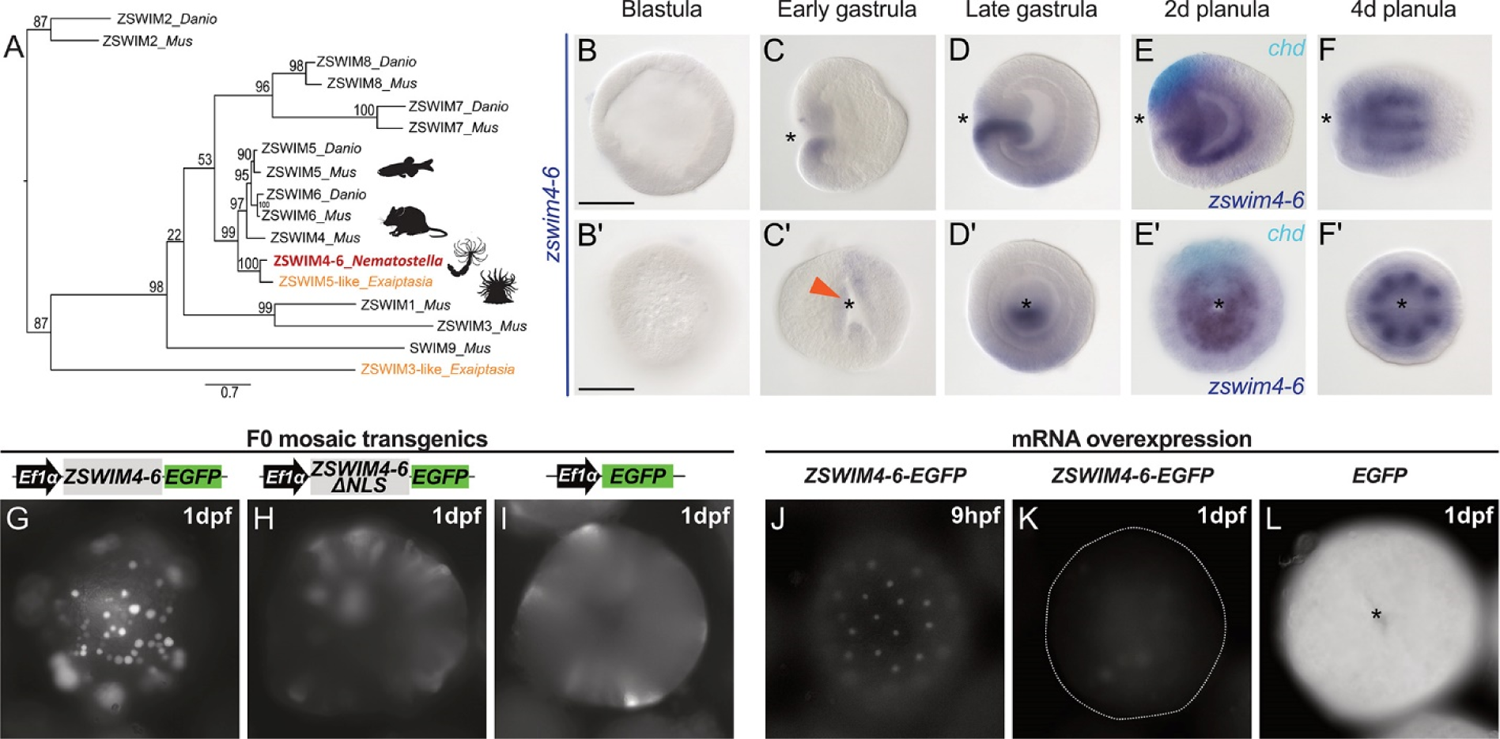
*zswim4-6* is a target of BMP signaling with bilaterally symmetric expression, and encodes a nuclear protein. (A) Maximum likelihood phylogeny shows that *Nematostella* ZSWIM4-6 clusters with ZSWIM4/5/6 from zebrafish and mouse. (B-F’) *Nematostella zswim4-6* expression follows the dynamic BMP signaling domain (see Fig. 1D for comparison). Double ISH shows *zswim4-6* and *chd* transcripts localize to the opposite sides of the directive axis. (G-I) Mosaic expression of ZSWIM4-6-EGFP under the control of the ubiquitously active *EF1α* promoter in F0 transgenic animals demonstrates that ZSWIM4-6 is a nuclear protein. Full-length ZSWIM4-6-EGFP is translocated into the nuclei (G), while ZSWIM4-6ΔNLS-EGFP missing the predicted NLS remains cytoplasmic (H), similar to the EGFP control (I). Exposure time was the same in all images. (J-L) Microinjection of *ZSWIM4-6-EGFP* mRNA results in a weak EGFP signal detectable in the nuclei of the early blastula (J), which progressively disappears towards late gastrula (K). EGFP translated from *EGFP* mRNA remains readily detectable (L). To visualize the weak signal in (J-K), the exposure had to be increased in comparison to (L). Asterisks mark the oral side; scale bars 100 µm.

Sequence analysis of the deduced ZSWIM4-6 protein revealed a potential N-terminal nuclear localization signal (NLS). To analyze the intracellular localization of ZSWIM4-6, we mosaically overexpressed an EGFP-tagged version of ZSWIM4-6 in F0 transgenic animals under the control of the ubiquitously active *EF1α* promoter (Kraus et al., 2016; Steinmetz et al., 2017). We compared the intracellular localization of the tagged wild type form to a truncated version of ZSWIM4-6 (ZSWIM4-6ΔNLS-EGFP), where the first 24 bases coding for the putative NLS were replaced by a start codon. Full-length ZSWIM4-6-EGFP was enriched in the cell nuclei (Fig. 4G), while the truncated version of ZSWIM4-6 lacking the NLS remained cytoplasmic, similar to the EGFP signal in transgenic animals overexpressing only cytoplasmic EGFP (Fig. 4H-I). Nuclear localization of ZSWIM4-6 could be confirmed by microinjection of *zswim4-6-EGFP* mRNA (Fig. 4J-L). Strikingly, the nuclear fluorescent signal of ZSWIM4-6-EGFP, which is weak but clearly visible at the blastula stage (Fig. 4J), progressively decreases and is barely detectable at 24 hpf (Fig. 4K). In contrast, injection of mRNA encoding cytoplasmic EGFP results in a strong fluorescent signal lasting for several days (Fig. 4L). Currently, the dynamics of the endogenous ZSWIM4-6 protein, as well as the mechanism behind this fast turnover after overexpression are unclear.

To analyze the function of BMP signaling in establishing the *zswim4-6* expression domain, we performed KDs of several components of the BMP signaling network (Fig. 5A). Previous studies have shown that the KDs of *bmp2/4* and *chd* suppress BMP signaling activity and abolish the pSMAD1/5 gradient (Genikhovich et al., 2015; Leclère and Rentzsch, 2014). A KD of another BMP ligand, GDF5-like, reduces BMP signaling activity, although a shallow BMP signaling gradient is preserved, while the knockdown of the BMP inhibitor Gremlin results in an expansion of the nuclear pSMAD1/5 gradient (Genikhovich et al., 2015). In line with the proposed role of BMP signaling in activating *zswim4-6* expression, the KDs of *bmp2/4* and *chd* led to a loss of *zswim4-6* expression (Fig. 5B-D’, Supplementary Data 1). In contrast, KD of *gdf5-like* and *gremlin* did not strongly affect *zswim4-6* expression (Fig. 5E-F’, Supplementary Data 1), suggesting that *zswim4-6* expression might either be robust towards milder alterations in pSMAD1/5 levels, or specifically depend on the BMP2/4-mediated, but not GDF5-like-mediated BMP signaling, or both.

**Figure 5.**
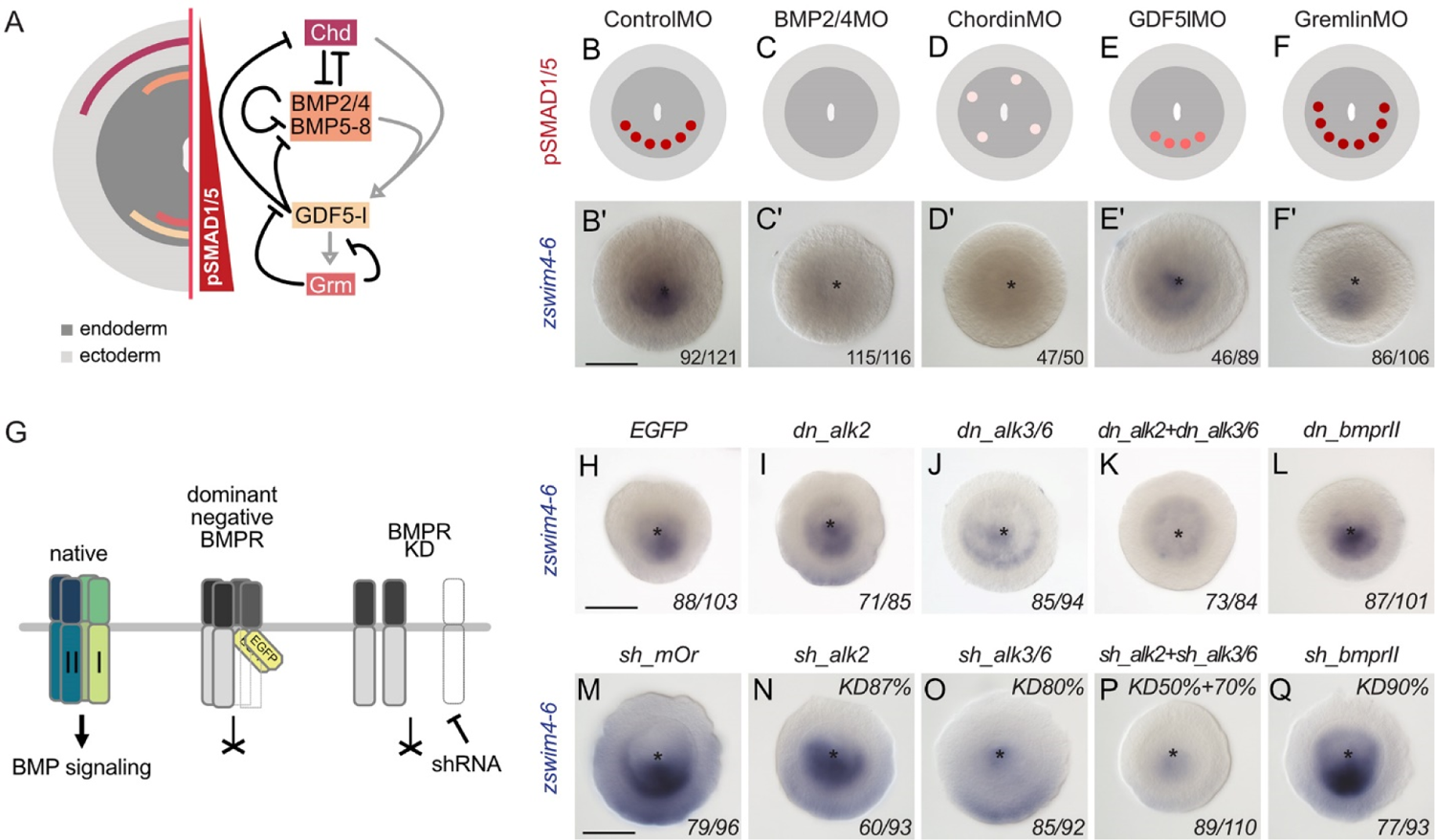
*zswim4-6* expression is regulated by BMP2/4 but not by GDF5-like signaling, and preferentially via the BMPRI receptor ALK3/6. (A) Expression domains and putative interactions of the BMP network components in 2d embryos (based on Genikhovich et al., 2015). (B-F’) Effect of the KD of the individual BMP signaling components on nuclear pSMAD1/5 and on *zswim4-6* expression. *zswim4-6* expression is abolished upon KD of *bmp2/4* and *chordin*, but not in KD of *gdf5-l* and *gremlin*. Sketches in B-F show that the gradient is lost upon BMP2/4 and Chd KD, reduced upon GDF5-like KD and expanded upon Gremlin KD (based on Genikhovich et al., 2015). (G) Overview of the expected effect of the dominant negative BMP receptor (Ser/Thr-kinase domain replaced with EGFP) overexpression and BMP receptor RNAi on BMP signaling. (H-L) *zswim4-6* expression is reduced upon overexpression of the dominant negative Type I BMP receptor dn_Alk3/6 and combined overexpression of dn_Alk2 and dn_Alk3/6 but remains unchanged in dn_Alk2 alone and dominant negative Type II BMP receptor dn_BMPRII. (M-Q) Expression of *zswim4-6* is reduced upon KD of the Type I BMP receptor Alk3/6 and combined KD of Alk2 and Alk3/6 but unaffected upon KD of Alk2 and BMPRII. Asterisks mark the oral side. Scale bars correspond to 100 µm. The numbers in the bottom right corner indicate the ratio of embryos displaying the phenotype shown in the image to the total number of embryos treated and stained as indicated in the figure.

To address these two possibilities in more detail, we first tested the response of *zswim4-6* to reduced levels of BMP signaling. We injected *Nematostella* embryos with different concentrations of BMP2/4MO starting with 50 µM and going up to the regular BMP2/4MO working concentration of 300 µM in steps of 50 µM, and stained these embryos with probes against *zswim4-6* and *bmp2/4*. Strikingly, within individual embryos, the response to the different concentrations of the BMP2/4MO was binary: either *zswim4-6* and *bmp2/4* expression appeared normal, or it was lost (*zswim4-6*) or radialized (*bmp2-4*), while intermediate phenotypes were never observed (Supplementary Fig. 2). However, the proportion of the embryos showing radialized bmp2/4 and abolished zswim4-6 expression increased depending on the amount of injected BMP2/4MO. This independently confirms an earlier observation of the binary effect of injecting iteratively decreasing concentrations of BMP2/4MO on mesentery formation (Leclère and Rentzsch, 2014). In their hands, the embryos either developed 8 or 0 mesenteries, but never some number in between, and the fraction of the embryos without mesenteries grew along with the increasing BMP2/4MO concentration. These results suggest that alterations in the BMP2/4-mediated signaling may not change the shape of the BMP signaling gradient, to which *zswim4-6* expression could be more or less robust, but rather define whether there is a BMP signaling gradient or not.

Since differential ligand selectivity of the BMP receptors was another possible explanation for the *zswim4-6* dependence on BMP2/4 but not on GDF5-like, we analyzed the expression of *zswim4-6* upon suppression of each of the two *Nematostella* BMP Type I receptors (*alk2* and *alk3/6*), and a single BMP Type II receptor (BMPRII) (Supplementary Fig. 3). We suppressed Alk2, Alk3/6 and BMPRII either by microinjecting mRNA of the dominant-negative version of each of these receptors, in which the intracellular kinase domain was replaced by EGFP, or by shRNA-mediated RNAi (Fig. 5G). In both assays, *zswim4-6* expression appears to be stronger affected by the Alk3/6 suppression than by the Alk2 suppression, and is nearly abolished by the combined Alk3/6/Alk2 KD (Fig. 5G-K, M-P). In contrast, it is largely unaffected by BMP Type II receptor suppression (Fig. 5G, L, Q). Thus, currently, our data favor the hypothesis that *zswim4-6* expression is regulated by BMP2/4-mediated signaling, which appears to be preferentially transduced by the Alk3/6 receptor or, possibly by the Alk3/6/Alk2 heterodimer, as it has been recently reported in fish (Tajer et al., 2021). Surprisingly, it also appears that the suppression of BMPRII during early *Nematostella* development can be compensated for - likely by another Type II TGFβ receptor molecule.

### ZSWIM4-6 is a modulator of BMP-dependent patterning

Given that *zswim4-6* expression is bilaterally symmetric and highly dependent on BMP signaling in *Nematostella*, we wanted to see whether *zswim4-6* itself contributes to the patterning of the directive axis. To this end, we performed KD of *zswim4-6* activity using a translation-blocking morpholino targeting *zswim4-6* (Supplementary Fig. 4).

Analysis of the overall morphology at late planula stage showed that *zswim4-6* KD leads to defects in endoderm compartmentalization. In control morphants, eight mesenteries reach all the way to the pharynx, and the primary polyp has four tentacles (Fig. 6A-A’’). In contrast, in ZSWIM4-6MO, the mesenterial chamber expressing HoxE protein fails to reach the pharynx resulting in the fusion of the two neighboring mesenterial segments (Fig. 6B-B’). Consequently, *zswim4-6* morphant primary polyps form only three instead of the typical four tentacles (Fig. 6B’’). Endoderm segmentation defects have previously been observed in *Nematostella* larvae with defects in pSMAD1/5 gradient formation, as well as in *Hox* gene activity (Genikhovich et al., 2015; He et al., 2018; Leclère and Rentzsch, 2014), and, based on its expression domain, *zswim4-6* may be involved in either of these two processes. However, in contrast to the *HoxE* mutants, which also have characteristic three-tentacle primary polyps due to the loss of the *HoxE*-expressing mesenterial chamber and the fusion of the *HoxD*-expressing mesenterial chambers capable of forming a tentacle (He et al., 2018), the affected pair of mesenteries is not lost in the *zswim4-6* morphants, and *HoxE* is detectable (Fig, 6B’). Therefore, we concentrated on analyzing the molecular role of *zswim4-6* in the BMP signaling network.

**Figure 6.**
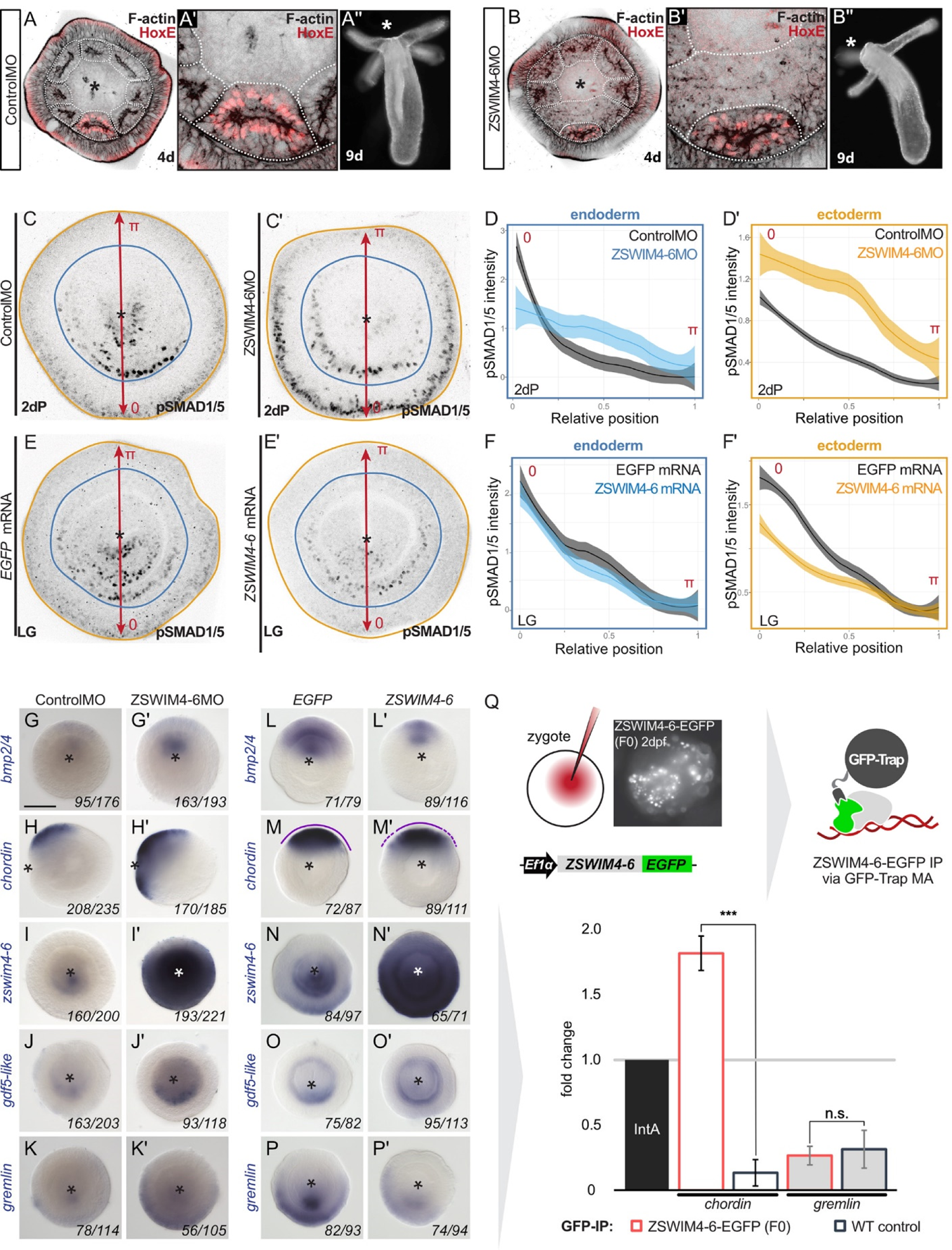
ZSWIM4-6 is a modulator of BMP signaling that appears to act as transcriptional repressor. (A-B) Morpholino KD of *zswim4-6* results in patterning defects. In the 4d planula, the HoxE-positive mesenterial chamber does not reach the pharynx, which leads to the fusion of neighboring chambers (compare A, A’ with B, B’). This results in the formation of three instead of four tentacles in the 9d polyp (compare A’’ with B’’). (C-F’) Immunofluorescence and quantification of relative nuclear anti-pSMAD1/5 staining intensities in 2d ZSWIM4-6 morphants (C-D’) and upon *zswim4-6* mRNA overexpression in the late gastrula (E-F’). Intensity measurements are plotted as a function of the relative position of each nucleus in the endoderm or in the ectoderm along a 180° arc from 0 (high signaling side) to π (low signaling side). The measurements from Control MO embryos (n=10) and ZSWIM4-6MO embryos (n=10), as well as *egfp* mRNA embryos (n=24) and *zswim4-6* mRNA embryos (n=8) are described by a LOESS smoothed curve (solid line) with a 99% confidence interval for the mean (shade). For visualization purposes, the intensity values were normalized to the upper quantile value among all replicates and conditions of each control-experiment pair. (G-K’) Expression of *zswim4-6* and BMP network components in the 2d planula upon morpholino KD of *zswim4-6*; All images except for H and H’ show oral views; (L-P’) Expression of *zswim4-6* and BMP network components in late gastrula (30 h) upon *zswim4-6* mRNA injection; oral views, purple dashed line marks the loss of a sharp boundary of *chd* expression. Asterisks mark the oral side. (Q) ChIP with GFP-Trap detects ZSWIM4-6-EGFP fusion protein in the vicinity of the pSMAD1/5 binding site in the upstream regulatory region of *chordin* but not of *gremlin*.

First, we assessed whether the pSMAD1/5 gradient is affected upon *zswim4-6* KD. To this end, we performed anti-pSMAD1/5 immunostainings in 2d planulae and quantified pSMAD1/5 levels along the directive axis. We found that the gradient profile of pSMAD1/5 in the endoderm was flattened in *zswim4-6* morphants compared to controls, and that peak levels of pSMAD1/5 activity were not reached. pSMAD1/5 activity in the ectoderm was, however, strongly elevated upon *zswim4-6* KD (Fig. 6C-C’, D-D’). To corroborate this, we then analyzed the pSMAD1/5 gradient in embryos microinjected with *zswim4-6-EGFP* mRNA. Due to the fast turnover of ZSWIM4-6-EGFP (see Fig. 4J-L), we quantified the gradient at the earliest possible stage after the symmetry break, i.e. at late gastrula. *zswim4-6* mRNA overexpression did not result in significant changes in the shape of the pSMAD1/5 gradient in the endoderm, however, ectodermal pSMAD1/5 levels were significantly reduced in the area where BMP signaling is normally strongest (Fig. 6E-E’, F-F’).

We then evaluated how KD of *zswim4-6* affected the transcription of markers expressed on the low BMP and high BMP signaling ends of the directive axis. First, we looked at *bmp2/4* and *chd* - the two key regulators of pSMAD1/5 gradient formation in *Nematostella*, whose expression is repressed by high levels of BMP signaling (Genikhovich et al., 2015; Leclère and Rentzsch, 2014; Saina et al., 2009). The *bmp2/4* and *chd* expression domains were expanded in *zswim4-6* morphants compared to controls (Fig. 6G-H’), which was especially evident in the ectodermal expansion of *chd*. Similarly, the *chd* expression domain was expanded in mosaic F0 *zswim4-6* knockouts generated by CRISPR/Cas9 to corroborate our morpholino KD results (Supplementary Fig. 5). This is striking as pSMAD1/5 stainings (Fig. 6C-D’) suggest that BMP signaling is stronger in the ectoderm of the morphants, therefore one could expect a reduction rather than an expansion of the *chordin* expression domain. Next, we looked at the markers of the high BMP signaling end: *zswim4-6*, *gdf5-like*, and *grm,* which are positively regulated by BMP signaling. The KD of *zswim4-6* translation resulted in a strong upregulation of *zswim4-6* transcription, especially in the ectoderm, suggesting that ZSWIM4-6 might either act as its own transcriptional repressor or that *zswim4-6* transcription reacts to the increased ectodermal pSMAD1/5 (Fig. 6I-I’). The expression of *gdf5-like* was largely unaffected (Fig. 6J-J’), while expression of *grm* was expanded in the ectoderm (Fig. 6K-K’), which reflects the flattening of the endodermal and the expansion of the ectodermal pSMAD1/5 gradient in the morphants (Fig. 6C-D’).

The overexpression of *zswim4-6-EGFP* by mRNA injection resulted in largely reciprocal effects on gene expression compared to morpholino-mediated KD, although less pronounced. As expected from the analysis of the morphants, *zswim4-6-egfp* overexpression resulted in a mild reduction of the expression domains of *bmp2/4* and *chd* (Fig. 6L-M’). Moreover, the otherwise very sharp border of the *chd* expression domain appeared diffuse (Fig. 6M’). *In situ* hybridization with the *zswim4-6* probe detected exogenous *zswim4-6* throughout the injected embryo (Fig. 6N-N’). Not much change was observed in the *gdf5-like* expression (Fig. 6O-O’), while *grm* was slightly reduced (Fig. 6P-P’), which also reflects the behavior of the pSMAD1/5 gradient (Fig. 6F-F’). Taken together, in the absence of ZSWIM4-6, changes in the expression of genes expressed on the high pSMAD1/5 side (*gdf5-like*, *grm*) appear to mimic the changes in the levels and range of the BMP signaling gradient. In contrast, genes repressed by BMP signaling and expressed on the low BMP signaling side of the directive axis (*bmp2/4*, *chd*) seem to expand their expression domains in the absence of ZSWIM4-6 despite the increase in pSMAD1/5 levels. One possible explanation for this observation is that BMP signaling-mediated gene repression is less effective in the absence of *ZSWIM4-6*.

If ZSWIM4-6 is indeed involved in the transcriptional repression of the pSMAD1/5 targets, we would expect to find ZSWIM4-6 bound to pSMAD1/5 binding sites at the genes repressed by BMP signaling, but not at the genes activated by BMP signaling. Due to the lack of an αZSWIM4-6 antibody and fast ZSWIM4-6-EGFP turnover in mRNA-injected embryos (Fig. 4J-K), we microinjected *EF1α::ZSWIM4-6-EGFP* plasmid (Fig. 4G) to perform αGFP-ChIP on mosaic F0 embryos, where *zswim4-6-egfp* is transcribed under control of a ubiquitously active promoter in a small fraction of cells in each embryo. Since the amount of chromatin that could be immunoprecipitated in such an experimental setup was very low, we were limited to selecting one negatively regulated direct BMP target and one positively regulated direct BMP target for the subsequent qPCR analysis. We chose pSMAD1/5 sites in the regulatory regions of the negatively regulated direct BMP target *chordin* and the positively regulated direct BMP target *gremlin*, and checked by qPCR whether they were bound by ZSWIM4-6-EGFP. Binding to the *Intergenic region IntA* (Schwaiger et al., 2021; Schwaiger et al., 2014) was used as a normalization control. The pSMAD1/5 binding site-containing region at the *gremlin* locus was similarly depleted in wild type and in *EF1α::ZSWIM4-6-EGFP* injected embryos upon αGFP-ChIP (0.35-fold enrichment and 0.27-fold enrichment respectively; insignificant according to the two-tailed t-test; P=0.496). On the other hand, the pSMAD1/5 binding site-containing region at the *chordin* locus was 1.82-fold enriched upon αGFP-ChIP in *EF1α::ZSWIM4-6-EGFP* embryos and 0.15-fold enriched (significant according to the two-tailed t-test; P=0.00032) in wild type embryos (Fig. 6Q). Thus, the enrichment of the pSMAD1/5 binding site-containing region at the *gremlin* locus in *EF1α::ZSWIM4-6-EGFP* embryos in comparison to the wild type embryos was 0.87-fold. In stark contrast, the enrichment of the pSMAD1/5 binding site-containing region at the *chordin* locus in *EF1α::ZSWIM4-6-EGFP* embryos in relation to wild type embryos was 12.13-fold. This suggests an interaction of ZSWIM4-6 with the pSMAD1/5 binding site of the *chordin* regulatory region, but not of the *gremlin* regulatory region, and supports the hypothesis that ZSWIM4-6 might act as a co-repressor for pSMAD1/5 targets.

Our comparison of evolutionary conserved pSMAD1/5 ChIP targets showed that the *Nematostella zswim4-6* homolog *zswim5* is a direct BMP target also in *Xenopus* (Fig. 3C), but its function as a modulator of vertebrate BMP signaling has not been addressed. To determine whether vertebrate ZSWIM5 also influences BMP-mediated patterning, we assessed the effect of *zswim5* overexpression in zebrafish embryos. Injection of increasing amounts of zebrafish *zswim5* mRNA into zebrafish embryos at the one-cell stage caused increasingly severe developmental defects (Fig. 7A-F”). These defects included shortened and twisted tails as well as disorganized head structures and partially resemble loss-of-function mutants of the BMP pathway, such as the *bmp2* mutant *swirl* (Kishimoto et al., 1997; Nguyen et al., 1998). *zswim5* overexpression also led to a dampening of the ventrally-peaking BMP signaling gradient in early gastrulation-stage zebrafish embryos, as detected by pSMAD1/5/9 immunofluorescence (Fig. 7G). The reduced amplitude of ventral BMP signaling caused by *zswim5* overexpression in zebrafish mirrors the decrease in pSMAD1/5 levels observed in the *Nematostella* ectoderm upon overexpression of *zswim4-6* (Fig. 6E-F’). Together, this suggests that the function of ZSWIM4-6 as a BMP signaling modulator has likely been conserved in evolution since before the cnidarian-bilaterian split.

**Figure 7.**
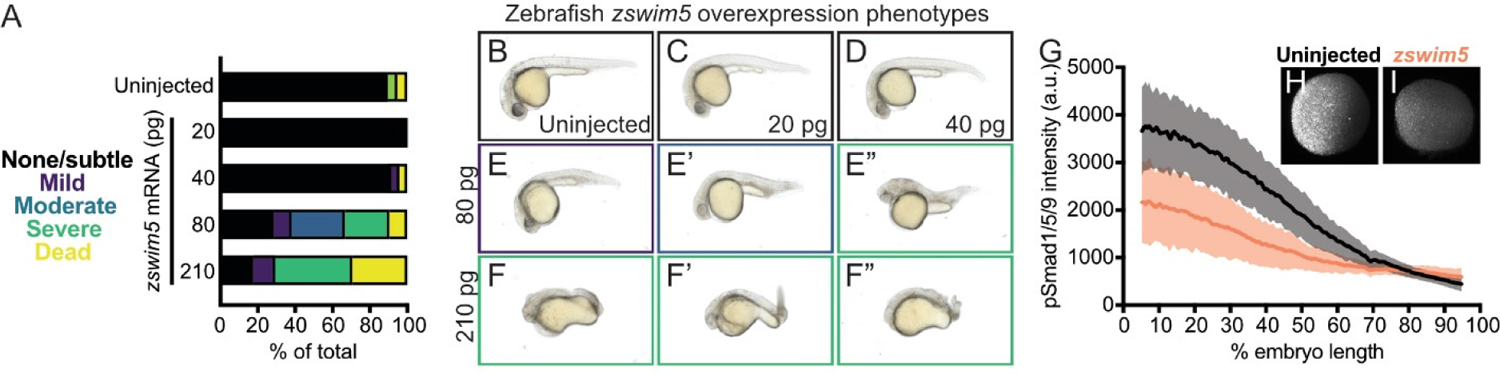
*Zswim5* overexpression dampens BMP signaling and causes developmental defects in zebrafish. (A-F”) Wild type zebrafish embryos were left uninjected (B) or injected at the one-cell stage with 20, 40, 80, or 210 pg *zswim5* mRNA. (A) Phenotype quantification at 1 day post-fertilization (dpf) shows increasingly severe developmental defects at higher amounts of mRNA. A representative selection was imaged at 1 dpf (B-F’’). Multiple embryos are shown for 80 (E-E”) and 210 (F-F”) pg to illustrate the variety of defects. Number of embryos - Uninjected: 19, 20 pg: 20, 40 pg: 24, 80 pg: 21, 210 pg: 17. (G-I) Embryos were injected with 80 pg *zswim5* mRNA or left uninjected and fixed at 50% epiboly (early gastrulation). BMP signaling levels were assessed using pSmad1/5/9 immunostaining. Animal pole views are shown with ventral on the left. (G) Quantification of immunofluorescence reveals lower amplitude BMP signaling gradients in *zswim5*-overexpressing embryos (I) compared to uninjected controls (H). Average background-subtracted intensity with standard deviation is plotted. Number of embryos - Uninjected: 8, *zswim5*-injected: 8.

## Discussion

### BMP signaling governs the expression of a second tier of developmental regulators

BMP signaling regulates the patterning of the anthozoan directive axis and the bilaterian D-V axis; however, the extent of conservation of downstream target genes between Bilateria and Cnidaria involved in the axial patterning has been largely unknown. In our αpSMAD1/5-ChIP-Seq in *Nematostella*, we identified putative direct targets of BMP signaling during the establishment of the directive axis in the late gastrula and in the fully compartmentalized 4d planula, providing new insights into the genetic program responsible for directive axis patterning. Functional annotation of the identified targets highlights an abundance of transcription factors (TFs) and signaling molecules (SMs) (Fig. 1F), suggesting that BMP signaling largely regulates a second tier of regulators rather than structural genes at the two stages we assayed. TFs and SMs were also prominent among the differentially expressed target genes upon knockdown of the BMP ligands BMP2/4 and GDF5-like (Fig. 1F), and among the targets with an identifiable BMP response element (BRE) (Katagiri et al., 2002). The presence of multiple members of Wnt, MAPK, Hedgehog, and Notch signaling pathways among the BMP signaling targets points at a high degree of coordination between different signal transduction cascades necessary to generate a properly organized embryo. Comparison of the *Nematostella* pSMAD1/5 targets with the available ChIP-seq data of the frog *Xenopus* (Stevens et al., 2017) and the fly *Drosophila* (Deignan et al., 2016), shows that BMP targets shared between Anthozoa and Bilateria are also mainly TFs and SMs.

Several molecules of the BMP pathway in Bilateria are known to be directly regulated by BMP signaling (Fig. 3C), (Deignan et al., 2016; Genander et al., 2014; Greenfeld et al., 2021; Rogers et al., 2020; Stevens et al., 2017). Likewise, pSMAD1/5 in *Nematostella* directly controls the transcription of multiple BMP network components (*chd, grm, bmprII, rgm, smad1/5, smad4*) and other associated regulators (*e2f4, morn, tob2, c-ski, tld1-like, bmp1-like, dusp1, dusp7*) that have not been characterized yet. Curiously, unlike *Xenopus bmp2, −4, −5, −7,* and *gdf2, −6, −9,* and *-10*, or *Drosophila dpp* (Deignan et al., 2016; Stevens et al., 2017), the promoters of *Nematostella* BMP-encoding genes do not appear to be directly bound by pSMAD1/5. Our ChIP-Seq data suggest that BMP signaling-dependent repression of *Nematostella bmp2/4* and *bmp5-8,* and BMP signaling-dependent activation of *gdf5-like* (Genikhovich et al., 2015) is indirect, but the link between the pSMAD1/5 activity and the transcriptional regulation of BMPs is still unknown.

### Staggered expression of Hox genes in *Nematostella* is directly controlled by BMP signaling

Bilateral body symmetry in *Nematostella* is manifested in the anatomy of the mesenteries subdividing the endoderm along the second body axis of the animal. In *Nematostella,* endodermal *hox* genes and *gbx* are expressed in staggered domains along the directive axis, and their expression boundaries exactly correspond to the positions of the emerging mesenteries (Ryan et al., 2007). This staggered endodermal *hox* and *gbx* expression is lost when BMP signaling is suppressed (Genikhovich et al., 2015). Recent loss-of-function analyses showed that RNAi of *hoxE*, *hoxD*, *hoxB* or *gbx* led to the loss of the pairs of mesenteries corresponding to the expression boundaries of the knocked down genes, while the knockdown of the putative co-factor of all Hox proteins, *Pbx*, resulted in the loss of all mesenteries (He et al., 2018) phenocopying the loss of BMP signaling (Genikhovich et al., 2015; Leclère and Rentzsch, 2014; Saina et al., 2009). Our ChIP-Seq analysis showed that the endodermal BMP-dependent staggered expression of *hox* genes and *gbx* is directly controlled by nuclear pSMAD1/5. In the future, it will be of great interest to test whether the loss of the mesenteries upon *BMP2/4*, *BMP5-8*, or *Chd* KD is only correlated with or actually caused by the resulting simultaneous suppression of all the staggered endodermal *hox* genes and *gbx* in the absence of BMP signaling (Genikhovich et al., 2015; He et al., 2018).

The question of whether Hox-dependent axial patterning was a feature of the cnidarian-bilaterian ancestor or evolved independently in Cnidaria and Bilateria remains debated. Different authors used different *hox* genes to homologize the different cnidarian body axes to the bilaterian ones (Arendt et al., 2016; DuBuc et al., 2018; Finnerty et al., 2004). The unclear orthology of cnidarian *hox* genes (Chourrout et al., 2006), as well as the staggered expression along a cnidarian body axis patterned by BMP signaling rather than along the one regulated by Wnt and FGF, as in Bilateria, allowed us to suggest that the involvement of staggered *hox* genes in axial patterning in anthozoans and bilaterians is probably convergent (Genikhovich et al., 2015; Genikhovich and Technau, 2017). This, however, does not exclude the possibility that regulation of some specific *hox* genes by BMP signaling was indeed conserved since before the cnidarian-bilaterian split. We find several *hox* genes as conserved direct BMP targets in *Xenopus* and *Drosophila*, and BMP-dependent regulation of *hox* expression has been reported in these models. For instance, in *Xenopus,* during the specification of the hindgut, direct pSMAD1/β-catenin interactions control the expression of *hoxa11*, *hoxb4* and *hoxd1* (Stevens et al., 2017), and in *Drosophila*, a BMP-Hox gene regulatory network mutually regulating *decapentaplegic*, *labial* and *deformed* was shown to be involved in the head morphogenesis (Stultz et al., 2012). Another indication in favor of the ancient origin of the BMP control over Hox-dependent processes is our finding that the gene encoding the TALE class homeodomain transcription factor Meis is among the ChIP targets conserved between *Nematostella*, *Xenopus* and *Drosophila*. Meis, together with another TALE class transcription factor Pbx, forms a trimeric complex essential for the Hox and Parahox function in the bilaterian anterior-posterior axis patterning (Merabet and Galliot, 2015). Hox proteins were also shown to directly interact with Meis/Pbx in *Nematostella* (Hudry et al., 2014). This interaction appears to be crucial for the function of *Nematostella* Hox proteins since *Pbx* KD resulted in the loss of all eight mesenteries (He et al., 2018), phenocopying the “no *hox*, no *gbx*” state of the endoderm with suppressed BMP signaling.

### Roles of BMP signaling during neurogenesis

In protostome and deuterostome Bilateria with centralized nervous systems, one of the key roles of BMP signaling is repression of neuroectoderm formation. In contrast, cnidarians possess diffuse nervous systems with certain local neural accumulations, which, however, cannot be considered ganglia or brains (Arendt et al., 2016; Kelava et al., 2015; Martin-Duran and Hejnol, 2021). There is no indication that the onset of nervous system development in *Nematostella* is affected by BMP signaling. The expression of neuronal terminal differentiation genes starts already at the blastula stage (Richards and Rentzsch, 2014), which is before the onset of detectable BMP signaling. During subsequent development, neurons continue to form in both germ layers in a radially symmetric manner (Nakanishi et al., 2012), regulated by Wnt, MAPK and Notch signaling (Layden et al., 2016; Layden and Martindale, 2014; Richards and Rentzsch, 2014; Richards and Rentzsch, 2015; Watanabe et al., 2014). The only known exception to this is the population of GLWamide+ neurons. They arise on the *bmp2/4*-expressing side of the directive axis (area of minimal BMP signaling) under control of the atonal-related protein Arp6 (neuroD), which is the only *Nematostella Arp* gene with a bilaterally symmetric expression (Watanabe et al., 2014). Our ChIP and RNA-Seq data showed that *Arp6* is directly suppressed by BMP. *Arp6*, however, is not the only *Nematostella* BMP target gene, whose bilaterian orthologs are implicated in the regulation of neuronal development. We also find *isl*, *irx*, *lmx, ashB, hmx3, atoh7,* and *soxC* - a *Sox4/Sox11* ortholog (the latter is also a BMP target in *Xenopus*) (Bergsland et al., 2006; Doucet-Beaupre et al., 2015; Liang et al., 2011; Miesfeld et al., 2020; Rodriguez-Seguel et al., 2009; Stevens et al., 2017; Tomita et al., 2000; Wang et al., 2004). Similarly, among the ChIP targets we find orthologs of the “canonical” bilaterian axon guidance molecules such as *rgm*, *ephrin B*, and *netrin* (Lai Wing Sun et al., 2011; Niederkofler et al., 2004; Williams et al., 2003); however, it remains unclear whether these molecules are involved in the regulation of neural development in *Nematostella* or have a different function. In Bilateria, these proteins are expressed in various non-neural contexts. For example, RGM was shown to act as a potentiator of BMP signaling in *Nematostella* and in Bilateria (Leclère and Rentzsch, 2014; Mueller, 2015), and Netrin was demonstrated to inhibit BMP signaling (Abdullah et al., 2021).

### *Nematostella* ZSWIM4-6 as a potential conveyor of the BMP signaling-mediated gene repression

One of the most enriched pSMAD1/5 ChIP-targets in *Nematostella* was *zswim4-6*. Members of the zinc-finger family with SWIM domain were found in Archaea, prokaryotes and eukaryotes, and they were suggested to be capable of interacting with other proteins or DNA (Makarova et al., 2002). *Nematostella zswim4-6* gene activity follows the dynamics of BMP signaling. ZSWIM4-6 is a nuclear protein, whose paralogs, ZSWIM4, -/5, and -/6 proteins are conserved in Bilateria. To-date, several studies have linked ZSWIM4/5/6 with vertebrate neurogenesis and forebrain development. For instance, in the mouse, the paralogues *zswim4-6* were found to be expressed in the forebrain with distinct spatiotemporal patterns (Chang et al., 2020; Chang et al., 2021). *Zswim6* knockout mice display abnormal striatal neuron morphology and motor function (Tischfield et al., 2017). Additionally, *zswim4* mutations frequently occur in patients with acute myelogenous leukemia (Walter et al., 2012), while *zswim6* mutations in mammals associate with acromelic frontonasal dysostosis (a rare disease characterized by craniofacial, brain and limb malformations), and schizophrenia (Smith et al., 2014; Tischfield et al., 2017). Interestingly, *zswim4/5* in the frog embryo and *zswim4/6* in transit-amplifying mouse hair follicle cells are also direct targets of BMP signaling (Genander et al., 2014; Stevens et al., 2017). This suggests that *zswim4*-6 is a conserved downstream target of BMP signaling and may contribute to BMP signaling-dependent patterning events in various contexts. However, the function of zswim4-6 homologs during early embryogenesis has remained largely unexplored.

Here we provide evidence that *Nematostella zswim*4-6 acts as a feedback regulator of the shape of the BMP signaling gradient, and its loss of function results in defects in tissue compartmentalization. While the extracellular signaling network setting up the pSMAD1/5 gradient along the directive axis in *Nematostella* has been addressed before (Genikhovich et al., 2015; Leclère and Rentzsch, 2014; Saina et al., 2009), our study provides the first example of a nuclear protein modulating the pSMAD1/5 gradient in a non-bilaterian organism. Strikingly, overexpression of a *Danio rerio zswim4-6* paralog, *zswim5*, significantly dampened pSMAD1/5 levels in zebrafish, and the embryos displayed dose-dependent developmental defects with increasing amounts of injected *zswim5* mRNA. This recapitulated the effect of *zswim4-6* overexpression on the pSMAD1/5 gradient in *Nematostella*, indicating that ZSWIM4-6 proteins may play a conserved role in modulating the pSMAD1/5 gradient in Bilateria and Anthozoa.

Analysis of the transcriptional response to *zswim4-6* KD showed that the genes known to be negatively regulated by BMP signaling were de-repressed in spite of the increased levels of nuclear pSMAD1/5, while the genes positively regulated by BMP signaling were barely affected. This allowed us to hypothesize that ZSWIM4-6 promotes BMP-mediated transcriptional repression, while positively regulated BMP targets react to changes in the pSMAD1/5 gradient and do not become repressed by ZSWIM4-6. In line with this, our ChIP analysis of EGFP-tagged *Nematostella* ZSWIM4-6 showed that ZSWIM4-6 can directly bind to the pSMAD1/5 binding site in the upstream regulatory region of at least one gene, *chordin*, which is directly repressed by BMP signaling, but not to the upstream regulatory region of *gremlin*, which, unlike *chordin*, is directly activated by BMP signaling. In the future, the generation of a tagged ZSWIM4-6 transgenic line followed by ChIP-Seq will clarify at the genome-wide scale whether or not ZSWIM4-6 directly acts exclusively on genes repressed by BMP signaling.

### *zswim4-6* is selectively activated by BMP2-4 but not by GDF5-like

KD of distinct *Nematostella* BMP ligands showed that not all perturbations lead to a similar reduction in the *zswim4-6* expression domain. While KD of BMP2/4 and its putative shuttle Chordin abolished *zswim4-6* expression, the effect of GDF5-like KD on *zswim4-6* was minimal, despite a striking reduction in the pSMAD1/5 levels caused by GDF5lMO injection (Genikhovich et al., 2015). We see two possible explanations for this phenomenon: i) it can reflect the different levels of reduction of pSMAD1/5 signaling intensity upon BMP2/4 and GDF5-like KD; ii) it can indicate different receptor or binding co-factor preferences for different BMP ligands. Our attempts to gradually reduce BMP signaling by injecting different concentrations of BMP2/4MO confirmed an earlier observation (Leclère and Rentzsch, 2014) that these perturbations result in binary response: the embryos appear either normal (normal pSMAD1/5 gradient and normal marker gene expression at early planula, 8 mesenteries at late planula), or completely radialized (abolished pSMAD1/5gradient, radialized *bmp2/4* expression, 0 mesenteries at late planula). In contrast, GDF5-like morphants display an intermediate phenotype with a shallow pSMAD1/5 gradient at early planula and 4 mesenteries at late planula (Genikhovich et al., 2015). This suggests that, in spite of converging at the point of SMAD1/5 phosphorylation, BMP2/4-mediated and GDF5-like-mediated signaling may differ by their mode of action, which can be achieved, for instance, by different BMP ligands having different receptor preferences.

*In vitro* and *in vivo* studies showed that in Bilateria, different BMPs elicit combinatorial, non-redundant responses (Chen et al., 2013; Klumpe et al., 2022) displaying complex, promiscuous binding to various BMP and Activin receptors (Klumpe et al., 2022; Mueller and Nickel, 2012; Nickel and Mueller, 2019; Su et al., 2022). In different contexts, different ligands show distinct receptor preferences. For example, the protein precursor of the *Drosophila* BMP5-8 paralog Glass bottom boat can be proteolytically processed into a long and a short isoform, which have different developmental functions and preferentially signal via different Type II BMP receptors (Anderson and Wharton, 2017). In mammalian cell culture and biosensor assays, BMP2 and BMP4 were shown to bind and signal via Type I BMPR receptors Alk3 and Alk6, while GDF5 preferentially bound Alk6 (Heinecke et al., 2009; Nickel and Mueller, 2019). Strikingly, GDF5 could be mutated to mimic the binding preferences of BMP2, however, the binding of GDF5 and BMP2 to the same set of receptors elicited different, and in some cases opposite, cellular responses, which suggests the presence of some yet unknown co-factors involved in signal transduction (Klammert et al., 2015). In contrast, in vivo, the mammalian BMP5-8 paralogs BMP6 and BMP7 seem to be able to signal only through Alk2, although they bind Alk3 with a 30-fold higher affinity in vitro (Nickel and Mueller, 2019). Recent evidence from zebrafish suggests that Alk3/6 and Alk2 (zebrafish has two *alk3*, two *alk6* and one *alk2* gene) are non-redundant. At physiological expression levels, BMP2 and BMP7 homodimers do not appear to signal at all during D-V patterning; however, BMP2/7 heterodimers signal very effectively and require both Alk3 and Alk2 Type I receptors for signaling (Tajer et al., 2021). In our hands, shRNA-mediated KD and overexpression of the dominant-negative forms of *Nematostella* Alk2 and Alk3/6 also suggested their non-redundant roles. *Zswim4-6* expression was not affected by Alk2 KD, but became weaker upon Alk3/6 KD and abolished upon simultaneous Alk2/Alk3/6 KD. In contrast, BMPR-II appeared to be expendable for this signaling interaction, suggesting that its loss of function might be compensated by a different Type II receptor. In summary, in *Nematostella*, BMP2/4-mediated signaling, but not GDF5-like-mediated signaling, regulates *zswim4-6* expression. It appears that this selective action of different BMP ligands takes place at the level of the ligand-receptor interaction. Our experiments suggest that BMP2/4 (or rather a putative BMP2/4/BMP5-8 heterodimer) requires Alk3/6 or a combination of Alk3/6 and Alk2 but not Alk2 alone to regulate *zswim4/6*.

## Conclusion

In this paper we present evidence showing that in the sea anemone *Nematostella vectensis*, BMP signaling directly controls the expression of a number of previously characterized and many novel regulators of their bilaterally symmetric body plan. We show that *gbx* and all the *hox* genes, previously shown to control the regionalization of the secondary body axis of *Nematostella,* are under direct BMP regulation. In our dataset, we identified BMP target genes conserved between Cnidaria and Bilateria, among which we found a novel modulator of BMP signaling, the putative transcriptional repressor *zswim4-6*, whose expression is tightly controlled by BMP2/4-but not by GDF5-like-mediated BMP signaling. Our paper provides a valuable resource for future studies of BMP signaling in Cnidaria and Bilateria and offers the first glimpse into the fascinating area of control of BMP ligand/receptor selectivity in a bilaterally symmetric non-bilaterian.

## Materials and Methods

### Animal husbandry and microinjection

Adult *Nematostella vectensis* polyps were kept in the dark at 18 °C in *Nematostella* medium (16‰ artificial seawater, NM) and induced for spawning in a 25 °C incubator with 10 h of constant illumination. Egg packages were fertilized for 30 min, de-jellied in 3% L-cysteine/NM and washed 6 times with NM. Microinjection was carried out under a Nikon TS100F Microscope, using an Eppendorf Femtojet and Narishige micromanipulators as described in (Renfer and Technau, 2017).

Adult TE zebrafish were kept in accordance with the guidelines of the State of Baden-Württemberg (Germany) and approved by the Regierungspräsidium Tübingen (35/9185.46–5, 35/9185.81–5). Zebrafish embryos were maintained at 28℃ in embryo medium (250 mg/l Instant Ocean salt in reverse osmosis water adjusted to pH 7 with NaHCO_3_). Microinjections were carried out using PV820 Pneumatic PicoPumps (World Precision Instruments), M-152 micromanipulators (Narishige) and 1B100f-4 capillaries (World Precision Instruments) shaped with P-1000 micropipette puller (Sutter Instrument Company).

### Transgenesis, gene overexpression and knockdown

The full-length sequence of *Nematostella zswim4-6* was isolated by RACE-PCR. Native *zswim4-6* (EF1α*::zswim4-6-egfp*) and a truncated version lacking the first 24 bases coding for the NLS (*EF1α::zswim4-6Δnls-egfp)* were cloned into *AscI* and *SbfI* sites in the *Nematostella* transgenesis vector, downstream of the ubiquitously active *EF1α* promoter, and *I-SceI* meganuclease-mediated transgenesis was performed as specified (Renfer and Technau, 2017). For mRNA synthesis, the *EF1α* promoter was removed by digestion with PacI and AscI, the ends were blunted, and the plasmid re-ligated, which placed *zswim4-6* directly downstream of the SP6 promoter. Capped mRNA was synthesized using an SP6 mMessage mMachine Transcription Kit (Life Technologies) and purified with the Invitrogen MEGAclear Transcription Clean-Up Kit (Ambion). For overexpression, 150ng/µl mRNA of *ZSWIM4-6-EGFP* or *EGFP* control were injected. *zswim4-6* knockout animals were generated using the IDT CRISPR/Cas9 (Alt-R CRISPR-Cas9 crRNA:tracrRNA) gRNA system. Genotyping of F0 mosaic mutants after *in situ* hybridization was performed as described in (Lebedeva et al., 2021). Target sequence for gRNAs and genotyping primers are listed in the Supplementary Table 1. Dominant-negative BMP receptors were generated by removing the C-terminal serine-threonine kinase domains of Alk2, Alk3/6 and BMPRII and replacing them with the *EGFP* coding sequence. mRNAs of dominant-negative receptors and *EGFP* control were injected at 60ng/µl (Supplementary Table 2).

The zebrafish *zswim5* cDNA was cloned into the pCS2+ vector as follows. RNA was obtained from 75% epiboly-stage embryos using TRIzol (Invitrogen) and reverse-transcribed into cDNA using SuperScript III First-Strand Synthesis SuperMix (Invitrogen). *zswim5* was then amplified from cDNA using the primers ATGGCGGAGGGACGTGGA and TTAACCGAAACGTTCCCGTACCA followed by the addition of ClaI and EcoRI restriction sites using the primers TTCTTTTTGCAGGATCCCATCGATGCCACCATGGCGGAGGGACGTGGA and TAGAGGCTCGAGAGGCCTTGAATTCTTAACCGAAACGTTCCCGTACCA. The resulting amplicon was cloned into pCS2+ using ClaI and EcoRI (NEB) digestion and Gibson assembly (NEB). To generate mRNA, the pCS2-zswim5 plasmid was linearized with NotI-HF (NEB) and purified using a Wizard SV Gel and PCR Clean-up System (Promega). mRNA was generated from linearized plasmid using an SP6 mMessage mMachine Transcription Kit (Thermo Fisher Scientific) and column-purified using an RNeasy Mini Kit (Qiagen). mRNA concentration was quantified using an Implen NanoPhotometer NP80. For phenotype assessment, TE wild-type zebrafish embryos were injected through the chorion at the one-cell stage with 20, 40, 80, or 210 pg *zswim5* mRNA. Unfertilized and damaged embryos were removed approximately 1.3 h post-injection, and embryos were incubated at 28°C. At 1 day post-fertilization, embryos were scored based on gross morphology visible through the chorion. Representative embryos were immobilized with MESAB, dechorionated manually, and imaged in 2% methylcellulose on an Axio Zoom V16 microscope (ZEISS).

KD of *bmp2/4*, *chordin*, *gremlin* and *gdf5-l* in *Nematostella* were performed using previously published morpholino oligonucleotides (Supplementary Tables 3-4) as described (Genikhovich et al., 2015; Saina et al., 2009). The activity of the new translation-blocking ZSWIM4/6MO (5’ CCGTCCATAGCTTGTACTGATCGAC) was tested by co-injecting morpholino with mRNA containing the *mCherry* coding sequence preceded by either wildtype or 5-mismatch ZSWIM4/6MO recognition sequence in frame with mCherry (Supplementary Fig. 4). Approximately 4pL (Renfer and Technau, 2017) of 500µM solutions of GrmMO and GDF5-lMO, and 300 µM solutions of BMP2/4MO, ChdMO and ZSWIM4/6MO were injected, unless indicated differently. shRNA-mediated BMP receptor KD were carried out as described (Karabulut et al., 2019) using injection or electroporation of 800ng/µl shRNA for *alk2*, *alk3/6* and *bmprII* (Supplementary Table 5). shRNA against *mOrange* was used as control in all KD. To determine KD specificity, two non-overlapping shRNAs were tested for each gene and efficiencies were estimated by *in situ* hybridization and qPCR (Supplementary Table 6). qPCR normalization was performed using primers against GADPH.

### Chromatin immunoprecipitation and RNA-Seq

Processing of *Nematostella* embryos of two developmental stages, namely late gastrula and late planula, for ChIP was performed as described previously (Kreslavsky et al., 2017; Schwaiger et al., 2014). Input DNA was taken aside after chromatin shearing and pre-blocking with Protein-G sepharose beads. Immunoprecipitation was performed overnight at 4°C using 300µg of chromatin per reaction and polyclonal rabbit anti-Phospho-Smad1 (Ser463/465)/Smad5 (Ser463/465)/Smad8 (Ser426/428) (Cell Signaling, #9511; 1:1000). Pulldown of the immunoprecipitated chromatin using protein-G sepharose beads and elution were performed as described (Schwaiger et al., 2014). Libraries for sequencing of the input and immunoprecipitated DNA were prepared using the NEBNext Ultra End Repair/dA-Tailing Module kit and NEBNextUltraLigation Module with subsequent PCR amplification using KAPA Real Time Library Amplification Kit. 50 bp PE Illumina HiSeq2500 sequencing was performed by the Vienna BioCenter Core Facilities.

Raw reads were trimmed using trimmomatic v0.32 (Bolger et al., 2014) using the paired end mode and the ILLUMINACLIP option, specifying TruSeq3-PE as the target adaptors, seed mismatches at 2, a palindrome clip threshold of 30 and a simple clip threshold of 10. Additionally, the leading and trailing quality threshold were 5, and reads under a length of 36 were filtered out. Trimmed reads were aligned to the *Nematostella vectensis* genome (Putnam et al., 2007) using the “mem” algorithm and default settings. Alignments were then for conversion to bedpe format using the samtools v1.3.1 (Li et al., 2009) name sort and fixmate utilities. Bedtools (Quinlan and Hall, 2010) was used to convert to bedpe. Peak calling was performed using a modified version of peakzilla (Bardet et al., 2011) (“peakzilla_qnorm_patched.py”) which used quantile normalization for peak height. Peaks were merged across lanes using a custom script (“join_peaks.py”) using a maximum distance of 100 bp. Joined peaks were then annotated using a custom script (“associate_genes.py”). Gene model assignment was then manually curated to discard wrong gene models. A filter was applied (“final_filter.py”) which required the peaks to have an enrichment score of at least 10 and have an overall score of at least 80. Motif analysis was done using the MEME-ChIP pipeline (Machanick and Bailey, 2011) with the anr model, a window range of 5-30, number of motifs 10, DREME e-value threshold of 0.05, centrimo score and evalue thresholds of 5 and 10, respectively, against the JASPAR 2018 core motif database. Custom scripts mentioned above can be found in the following github repository: https://github.com/nijibabulu/chip_tools.

For RNA-seq, total RNA was extracted from 2 dpf embryos with TRIZOL (Invitrogen) according to the manufacturer’s protocol. Poly-A-enriched mRNA library preparation (Lexogen) and 50 bp SE Illumina HiSeq2500 sequencing was performed by the Vienna BioCenter Core Facilities. The number of the sequenced biological replicates is shown in the Supplementary Fig. 1. The reads were aligned with STAR (Dobin et al., 2013) to the *Nematostella vectensis* genome (Putnam et al., 2007) using the ENCODE standard options, with the exception that --alignIntronMax was set to 100 kb. Hits to the NVE gene models v2.0 (https://figshare.com/articles/Nematostella_vectensis_transcriptome_and_gene_models_v2_0/8 07696), were quantified with featureCounts (Liao et al., 2014), and differential expression analysis was performed with DeSeq2 (Love et al., 2014). Expression changes in genes with the adjusted p-value < 0.05 were considered significant. No additional expression fold change cutoff was imposed. Putative transcription factors (TF) and secreted molecules (SignalP) were identified using INTERPROSCAN (Madeira et al., 2019).

### Low input ChIP

EGFP-IP was performed on *EF1α::zswim4-6-egfp* F0 transgenic *Nematostella* 30 hpf. 4000-6000 embryos were used per replicate. Embryos were washed 2x with NM, 2x with ice-cold 1xPBS and collected in a 2 ml tube. They were fixed in 1 part of 50 mM HEPES pH 8, 1 mM EDTA pH 8, 0.5 mM EGTA pH 8, 100 mM NaCl and 3 parts of 1x PBS and 2% methanol-free formaldehyde (Sigma) for 15 min at RT on an overhead rotator. The fixative was removed and exchanged with 125 mM glycine in PBS for 10 min and the embryos were washed 3x with PBS. For long-term storage, embryos were equilibrated in HEG buffer (50 mM HEPES pH 7.5, 1 mM EDTA, 20% glycerol), as much liquid as possible was removed, and embryos were snap-frozen in liquid nitrogen and stored at –80°. Before sonication, embryos were thawed on ice and resuspended in 1ml E1 buffer (50 mM HEPES pH 7.5, 140 mM NaCl, 1 mM EDTA, 10% glycerol, 0.5% NP40, 0.25% Triton X-100, 1 mM PMSF, 1x Complete Protease Inhibitor (Roche), 1 mM DTT) and homogenized in a 1 ml dounce homogenizer with a tight pestle for 5 min directly on ice. The homogenate was centrifuged twice for 10 min at 4°C 1500 g, the supernatant was discarded and both pellets were resuspended in 130 µl lysis buffer (50 mM HEPES pH 7.5, 500 mM NaCl, 1 mM EDTA, 0.1% SDS, 0.5% N-Laurylsarcosine sodium, 0.3% Triton X-100, 0.1% sodium deoxycholate, 1 mM PMSF and 1x Complete Protease Inhibitor) and incubated in lysis buffer for 30min. The samples were sonicated in 130µl glass tubes in a Covaris S220 with following settings: for 20 s repeated 4 times: peak power=175, duty factor=20, cycles/burts=200; with pausing steps (peak power=2.5, duty factor=0.1, cycles/burts=5;) for 9 s in between each treatment.

After sonication, the sample was transferred to a low-bind Eppendorf tube and centrifuged at 4°C for 10 min at 16000 g. For the input fraction, 15 µl of the sample were taken aside and stored at −20°C. For the IP, the rest of the sample was filled up to 1ml with dilution buffer (10 mM Tris/HCl pH 7.5; 150 mM NaCl; 0.5 mM EDTA). GFP-Trap MA beads (Chromotek) were prepared according to the manufacturer’s instructions and incubated with the sample at 4°C overnight under rotation. The samples were washed 1x with dilution buffer, 1x with high salt dilution buffer (500 mM NaCl), again with dilution buffer and eluted twice with 50 µl 0.2 M glycine pH 2.5 within 5 min under pipetting. The eluates were immediately neutralized using 20 µl 1 M Tris pH 8. The IP fractions were filled up to 200 µl with MiliQ water, and the input fractions were filled up to 200µl with dilution buffer and incubated with 4µl of 10 mg/ml RNase A for 30 min at 37°C. Then 10 mM EDTA pH 8.0, 180 mM NaCl and 100µg/ml Proteinase K were added and incubated overnight at 65°C. The samples were purified using a Qiagen PCR Purification Kit. The IP fraction was eluted in 40 µl elution buffer, and the input fraction was eluted in 100 µl. The regulatory regions identified by pSMAD1/5 ChIP-Seq of *chordin* and *gremlin* were tested for enrichment by qPCR. Primers against the intergenic region A (*IntA*, scaffold_96:609598-609697) were used for normalization.

### Orthology analysis

The *Nematostella* proteome was downloaded from a public repository https://doi.org/10.6084/m9.figshare.807696.v2. The proteome from *Drosophila melanogaster* version BDGP6, corresponding to that used in Deignan et al. 2016, was downloaded from FlyBase (Larkin et al., 2021). The *Xenopus laevis* proteome version 9.1, corresponding to that used in (Stevens et al., 2017), was downloaded from the JGI genome browser (Nordberg et al., 2014). Orthology links between all proteomes were inferred using NCBI blastp (Altschul et al., 1990), requiring an e-value of at most 1e-5. Reciprocal best blast hits were determined using the bit score as the metric. For *X. laevis*, the genome suffix (.S or .L) was ignored, and only genes with BLAST hits in all species were considered (Fig. 3). The resulting hits were filtered for genes found as pSMAD1/5 targets in this study and in *Drosophila* and *Xenopus* analyses (Deignan et al., 2016; Stevens et al., 2017).

### *In situ* hybridization

Fragments for preparing in situ probes were amplified by PCR on ss cDNA (Supplementary Table 7) and cloned. *In situ* hybridization was carried out as described previously (Lebedeva et al., 2021), with minor changes. *Nematostella* embryos were fixed for 1 h in 4% PFA/PTW (1x PBS, 0.1% Tween 20) at room temperature. Permeabilization was performed for 20 min in 10 μg/ml Proteinase K/PTW. 4d planulae were incubated for 40min in 1U/μl RNAseT1 in 2xSSC at 37°C after the initial 2xSSC wash and before the 0.075xSSC washes.

### Phylogenetic analysis

Protein sequences of ZSWIM4-6 and related proteins (Supplementary Data 2) were aligned with MUSCLE using MegaX (Kumar et al., 2018) and trimmed with TrimAl 1.3 using an Automated 1 setting (Capella-Gutierrez et al., 2009). A maximum likelihood tree (JTTDCMut+F+G4, bootstrap 100) was calculated using IQTREE (Trifinopoulos et al., 2016).

### Antibody staining and pSMAD1/5 quantification

Immunostainings in *Nematostella* were performed according to Leclère and Rentzsch (2014) with minor changes. Embryos were fixed for 2min in ice-cold 2.5% glutaraldehyde/3.7% formaldehyde/PTX (1x PBS, 0.3% Triton-X), then transferred to ice-cold 3.7% FA/PTX and fixed at 4°C for 1 h. The samples were washed 5x with PTX, incubated in ice-cold methanol for 8 min, and washed 3x with PTX. After 2 h in the blocking solution containing 5% heat-inactivated sheep serum/95% PTX-BSA (PTX-BSA = 1% BSA/PTX), the samples were incubated overnight with the primary antibody (rabbit αpSMAD1/5/9 (Cell Signaling, #13820; 1:200) or rat αNvHoxE antibody (1:500)) in blocking solution. Embryos were then washed 5x with PTX, blocked for 2 h in blocking solution and incubated overnight with secondary antibody (goat α-rabbit IgG-Alexa568 or goat α-rabbit IgG-Alexa633) and DAPI (1:1000) in blocking solution. After the secondary antibody staining, the samples were washed 5x with PTX, mounted in Vectashield (VectorLabs), and imaged under Leica SP5X or Leica SP8 confocal microscopes. pSMAD1/5 staining intensities were quantified as described in Genikhovich et al., 2015. Control MO and ZSWIM4-6MO embryos, as well as *zswim4-6* mRNA and *egfp* injected embryos were stained with αpSMAD1/5 and DAPI and 16-bit images of single confocal sections of oral views were collected. The intensities of the pSMAD1/5 staining of the nuclei were measured in a 180° arc of from 0 to π using Fiji (Schindelin et al., 2012). Measurements in the endodermal and the ectodermal body wall were performed separately. The nuclei were selected in the DAPI channel and measured in the pSMAD1/5 channel, starting in the middle of the domain where the pSMAD1/5 signal is strongest and moving towards the position on the opposite side of the embryo. Every nucleus in each embryo was assigned a coordinate along the 0 to π arc by dividing the sequential number of each nucleus by the total number of analyzed nuclei in that particular embryo. αpSMAD1/5 staining intensities were then depicted as a function of the relative position of the nucleus.

TE wild type zebrafish embryos were dechorionated using Pronase and injected at the one-cell stage with 20-210 pg *zswim5* mRNA (80 pg mRNA was injected for the quantification). Unfertilized and damaged embryos were removed ∼1.5 h post-injection, and embryos were incubated at 28°C. At 50% epiboly embryos were fixed in cold 4% formaldehyde diluted in PBS. Uninjected embryos reached 50% epiboly 30-45 min before injected siblings. Fixed embryos were stored at 4°C overnight, washed with PBST, transferred to methanol (three PBST washes followed by three methanol washes), and stored at −20°C for at least 2 h. Embryos were then washed 4-5 times with PBST, blocked in FBS-PBST (10% FBS, 1% DMSO, 0.1% Tween-20 in PBS) for at least 1 h and then incubated in 1:100 anti-pSmad1/5/9 (Cell Signaling, #13820) in FBS-PBST at 4°C overnight. After 5-7 PBST washes over a period of 2-6 h, embryos were incubated again in FBS-PBST for at least 1 h and then transferred to 1:500 goat-anti-rabbit-Alexa488 (Life Technologies, A11008) and 1:1000 DRAQ7 (Invitrogen, D15105) overnight at 4°C. Embryos were washed 5-7 times with PBST over a period of 2-6 h and stored at 4°C overnight. pSmad1/5/9 immunofluorescence and DRAQ7 nuclear signal were imaged on a Lightsheet Z.1 microscope (ZEISS). Briefly, embryos were mounted in 1% low melting point agarose and imaged in water using a 20× objective. Embryos were oriented with the animal pole toward the imaging objective, and 50-90 z-slices with 7 μm between each slice were acquired. Quantification was performed on maximum intensity projections. Images were manually rotated in Fiji with the ventral side to the left, an ROI was manually drawn around each embryo to exclude non-embryo background, and the ventral-to-dorsal intensity profile was determined by calculating the average intensity in each column of pixels (pixel size: 0.457 μm × 0.457 μm). The profiles were binned and normalized to % length to account for embryo-to-embryo variability in length (Rogers et al., 2020). For background subtraction, intensities measured as described above in four randomly oriented controls not exposed to the primary antibody were subtracted from intensity profiles.

## Acknowledgements

This work was funded by the Austrian Science Foundation (FWF) grants P26962-B21 and P32705-B to G.G. and by the European Research Council (ERC) grants 637840 (*QUANTPATTERN*) and 863952 (*ACE-OF-SPACE*) to P.M.. We thank Michaela Schwaiger, Taras Kreslavsky, Hiromi Tagoh and Patricio Ferrer Murguia for their help with the ChIP protocol, Matthias Richter and Christian Hofer for their assistance with *in situ* analyses, Catrin Weiler for the assistance in cloning zebrafish *zswim5*, David Mörsdorf for critically reading the manuscript, and the Core Facility for Cell Imaging and Ultrastructure Research of the University of Vienna for access to the confocal microscope.

## Author contributions

A.S. and P.K. performed the majority of *Nematostella* experiments, analyzed data and wrote the paper. B.Z. performed the bioinformatic analyses. P.M. designed and supervised the zebrafish experiments. A.P. carried out pilot zebrafish experiments, and K.W.R. generated the data and analyses shown in Fig. 7. G.G. conceived the study, performed experiments, analyzed data and wrote the paper. All authors edited the paper.

## Data availability

All the sequencing data are available at NCBI as an SRA project PRJNA820593.

## Competing interests

The authors declare no competing interests

## Supplemental Information

**Supplementary video 1 –** At the 4d planula larva stage, BMP signaling activity locates to the mesenteries and in the ectoderm to eight adjacent stripes that merge in pairs between the future tentacle buds to form a circumoral ring.

**Supplementary data 1 –** anti-pSMAD1/5 ChIP analysis and RNA-Seq analysis of BMP2/4 and GDF5-l morphants.

**Supplementary data 2 –** Trimmed alignment of ZSWIM proteins

**Supplementary Fig 1.**
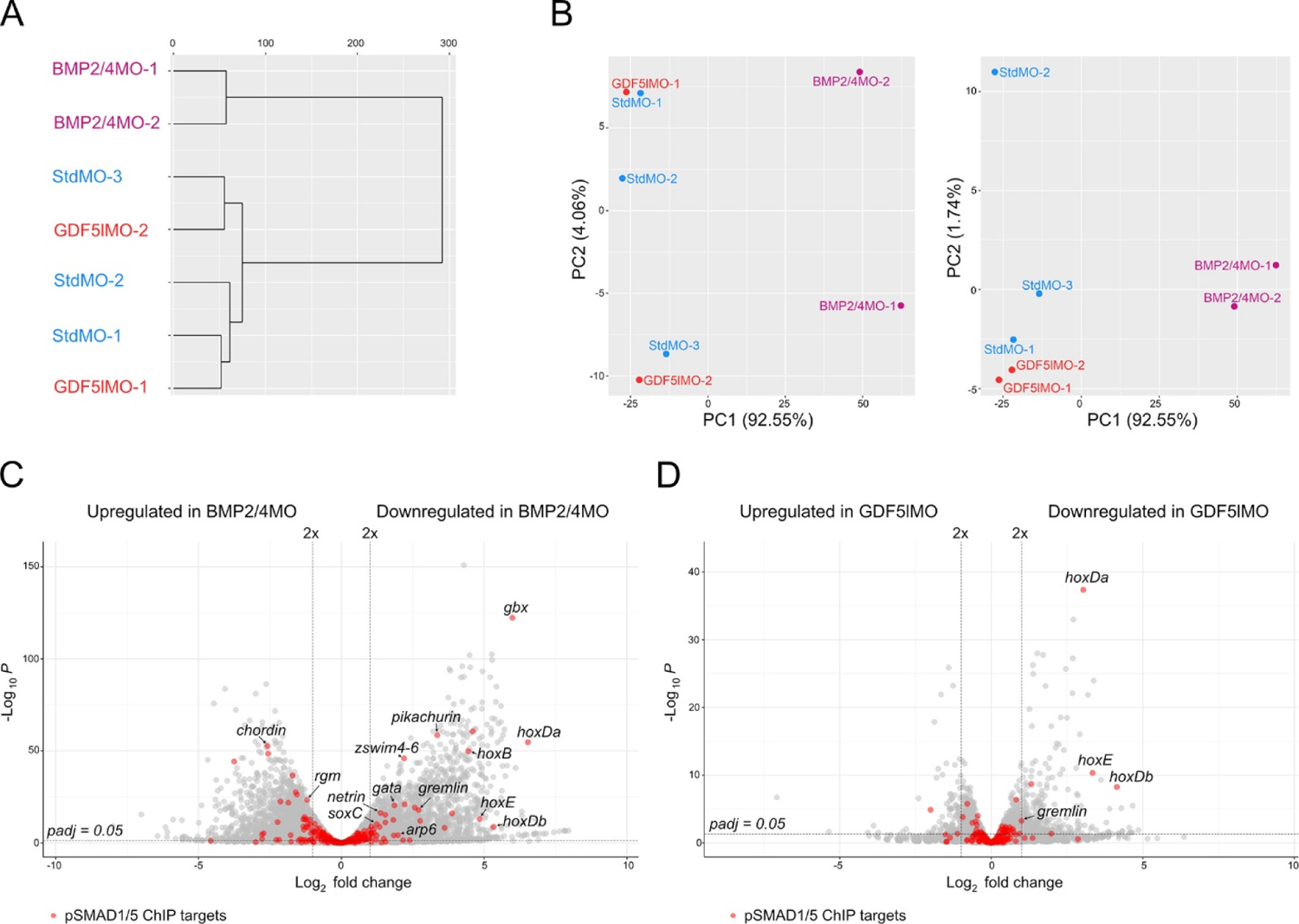
Transcriptomic comparison of BMP2/4, GDF5-l and control morphants and differential expression of pSMAD1/5 ChIP targets upon different knockdowns. (A) In the cluster dendrogram and (B) principal component analysis, replicates of GDF5–lMO (red) and StdMO (blue) transcriptomes group together, while the transcriptome of BMP2/4MO (magenta) is separated. (C-D) Volcano plots highlight differentially expressed pSMAD1/5 ChIP targets (red) in (C) BMP2/4 morphants and (D) GDF5-l morphants (p_adj_ ≤ 0.05).

**Supplementary Fig. 2.**
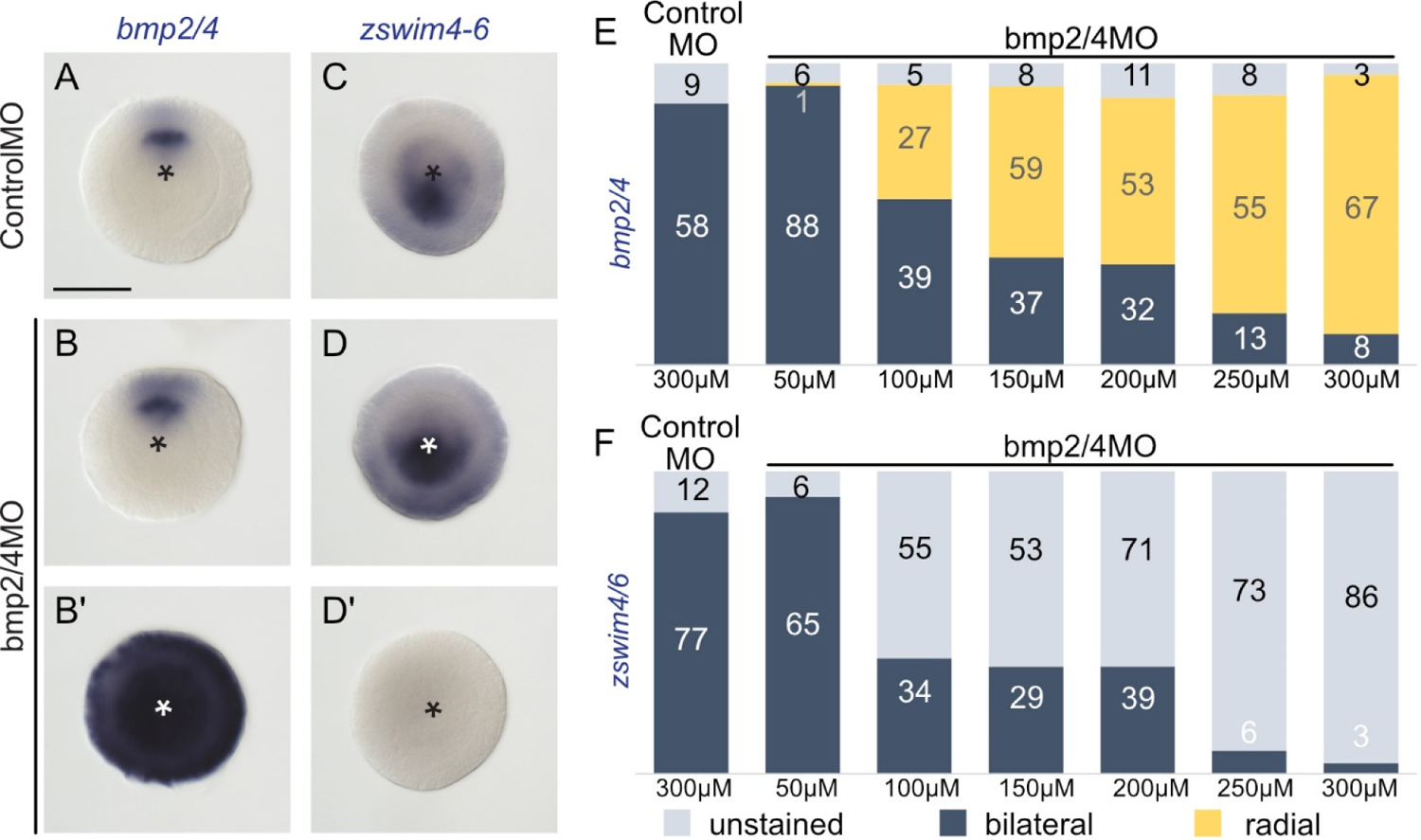
Knockdown of *bmp2/4* using different concentrations of bmp2/4 morpholino results in either normal, bilaterally symmetric expression or complete radialization but no intermediate phenotypes. (A) In the 2d planula normal bmp2/4 expression is confined to a narrow stripe on one side of the directive axis. Injection of different concentrations of bmp2/4MO results in either (B) no change of the normal expression pattern or (B’) complete radialization. (C) *zswim4/6* is expressed asymmetrically, on the opposite side of the directive axis. Different concentrations of bmp2/4MO result in either (D) normal *zswim4-6* expression or (D’) suppression of *zswim4-6*. (E) With the injection of increasing concentrations of *bmp2/4* morpholino, the number of embryos displaying a complete radialization of *bmp2/4* expression increases and (F) the number of embryos expressing *zswim4-6* decreases.

**Supplementary Fig. 3.**
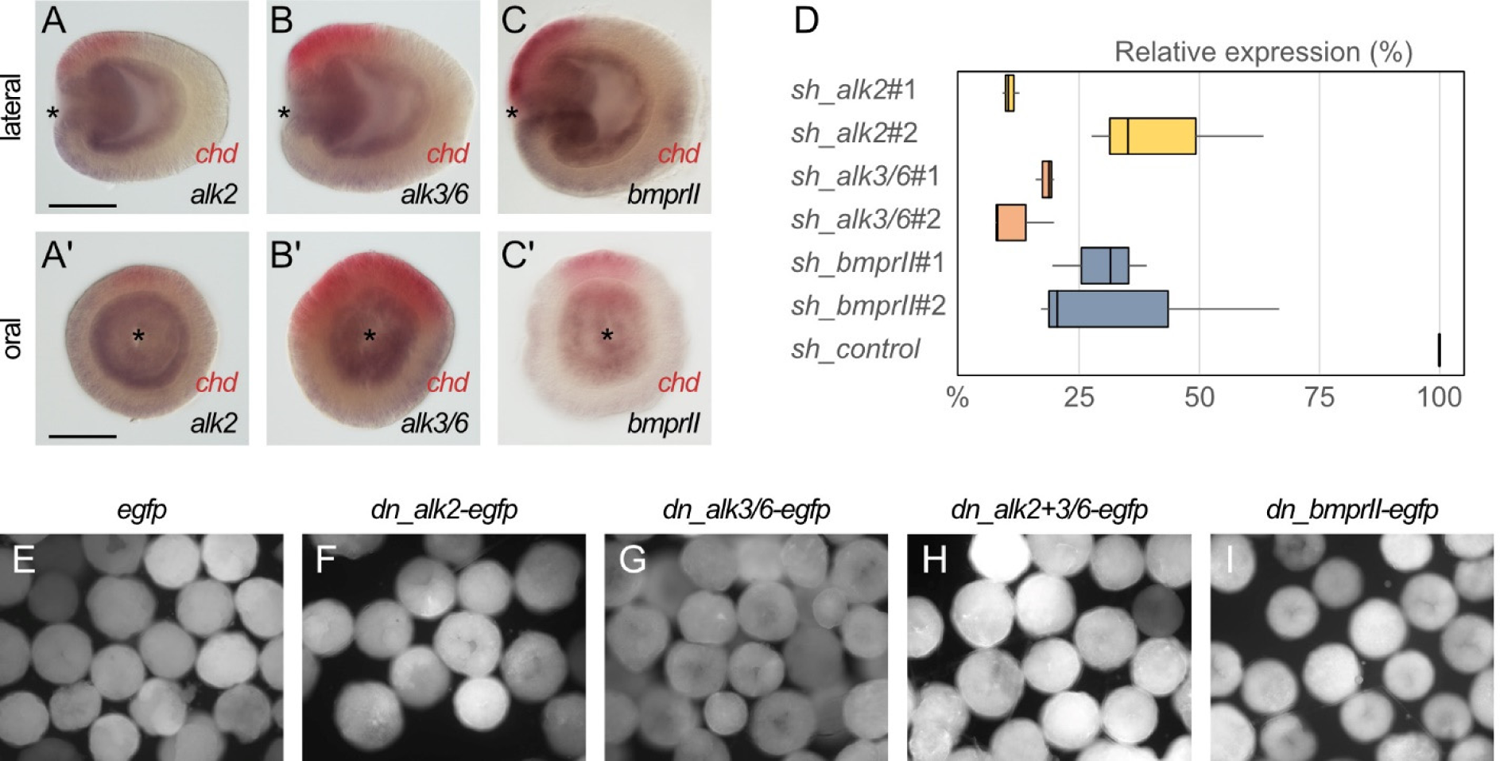
Expression of BMP receptors, efficiencies of RNAi mediated knockdowns and overexpression of dominant negative (dn) receptor constructs. Expression of BMP receptors (A-A’) *alk2*, (B-B’) *alk3/6* and (C-C’) *bmprII* is broadly detectable in endoderm with stronger expression on the high-BMP signaling activity side of the directive axis, opposite to *chordin* (red). (D) qPCR quantification of KD efficiencies for BMP receptor shRNAs. (E-I) EGFP signal in late gastrula embryos injected with (E) *egfp* mRNA and (F-I) mRNA of dominant-negative BMP receptor constructs. Asterisks mark the oral side, scale bars correspond to 100 µm.

**Supplementary Fig. 4.**
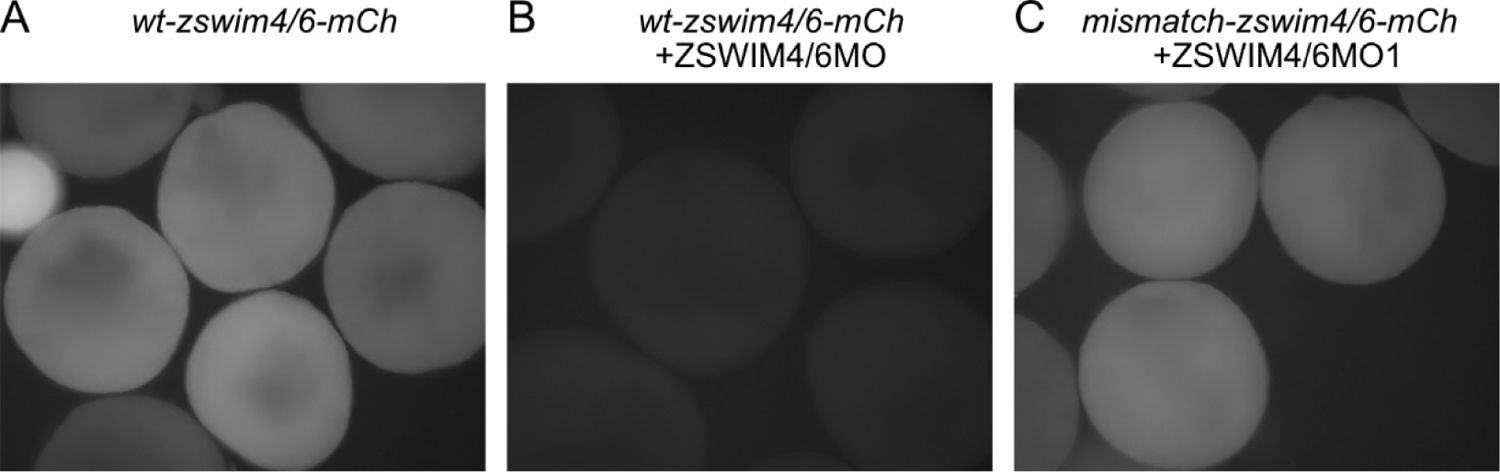
Testing ZSWIM4/6 morpholino specificity. (A) Fluorescent signal in embryos injected with mRNA coding for the ZSWIM4/6 recognition sequence fused to the mCherry (*wt-zswim4/6-mCh*) coding sequence. (B) Co-injection with ZSWIM4/6 morpholino can suppress the translation of mRNAs containing the respective recognition sequence. (C) Translation of mRNA coding for the zswim4/6 recognition sequence carrying 5 mismatches and fused to mCherry (*mismatch-zswim4/6-mCh*) is no longer suppressed when co-injected with ZSWIM4/6 morpholino.

**Supplementary Fig. 5.**
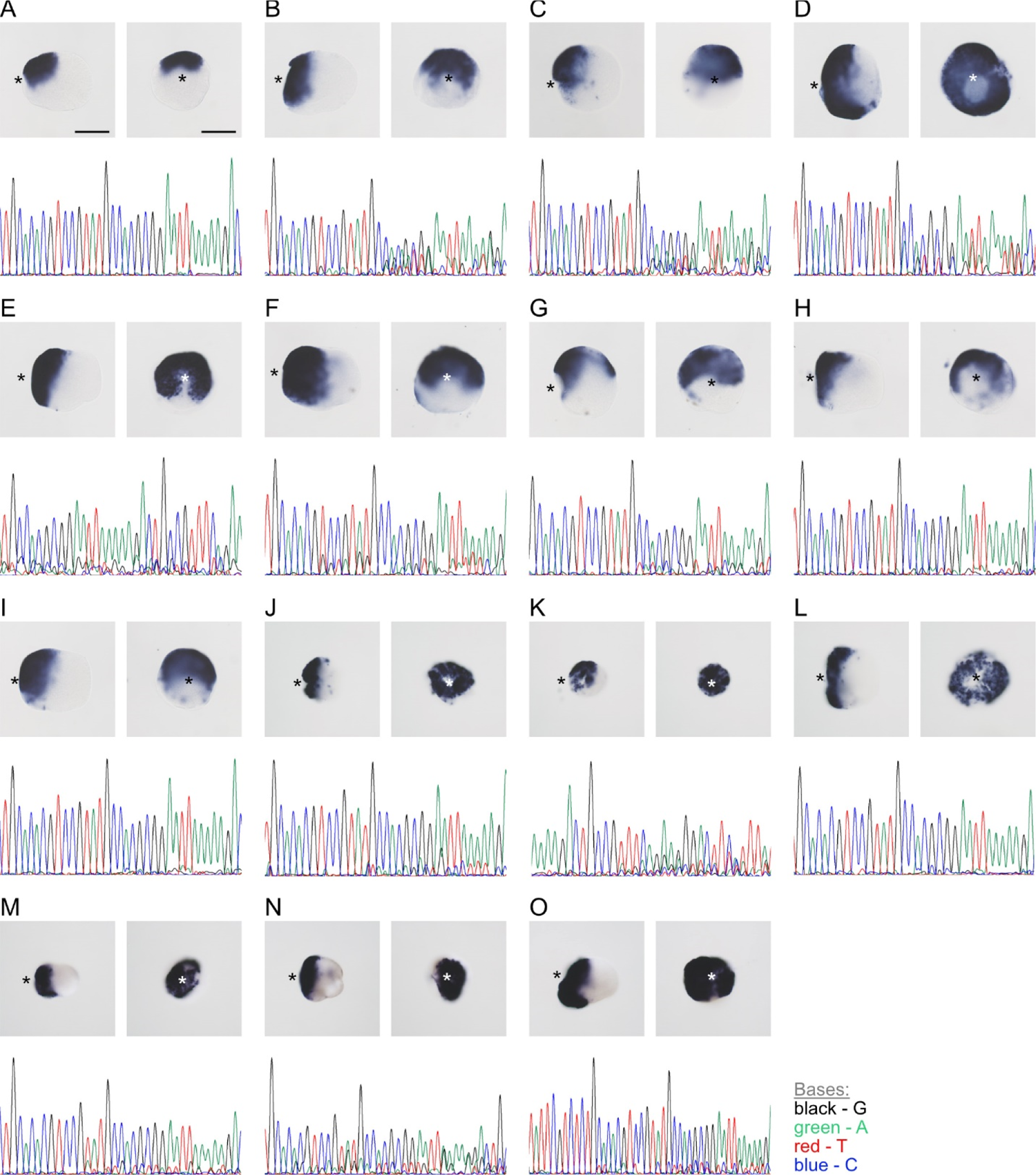
CRISPR/Cas9 mediated mutagenesis of *zswim4-6* results in the expansion of *chordin* expression in mosaic mutants (F0). (A) Bilateral expression of *chordin* in wild type 2d planula and corresponding sequencing chromatogram of *zswim4-6* guide RNA target region. (B-O) expanded or radialized chordin expression in 2d mutant planulae injected with a single guide RNA (B-I) or two guide RNAs (J-O) targeting the SWIM zinc-finger domain. Corresponding sequencing chromatograms show variability in the target loci of mutant animals. Asterisks mark the oral side, scale bars represent 100µm.

---

**Supplementary Table 1.**
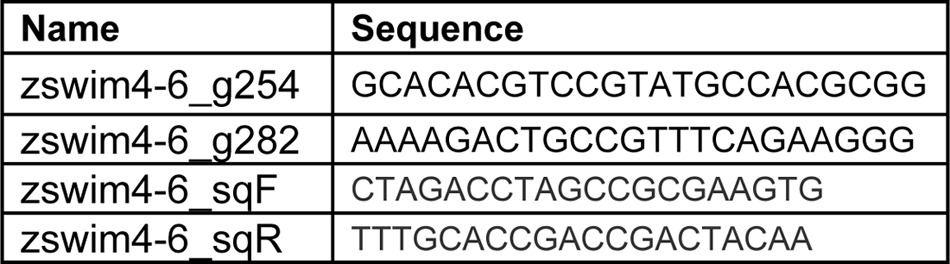
ZSWIM4-6 guide RNA sequences and sequencing primers

**Supplementary Table 2.**
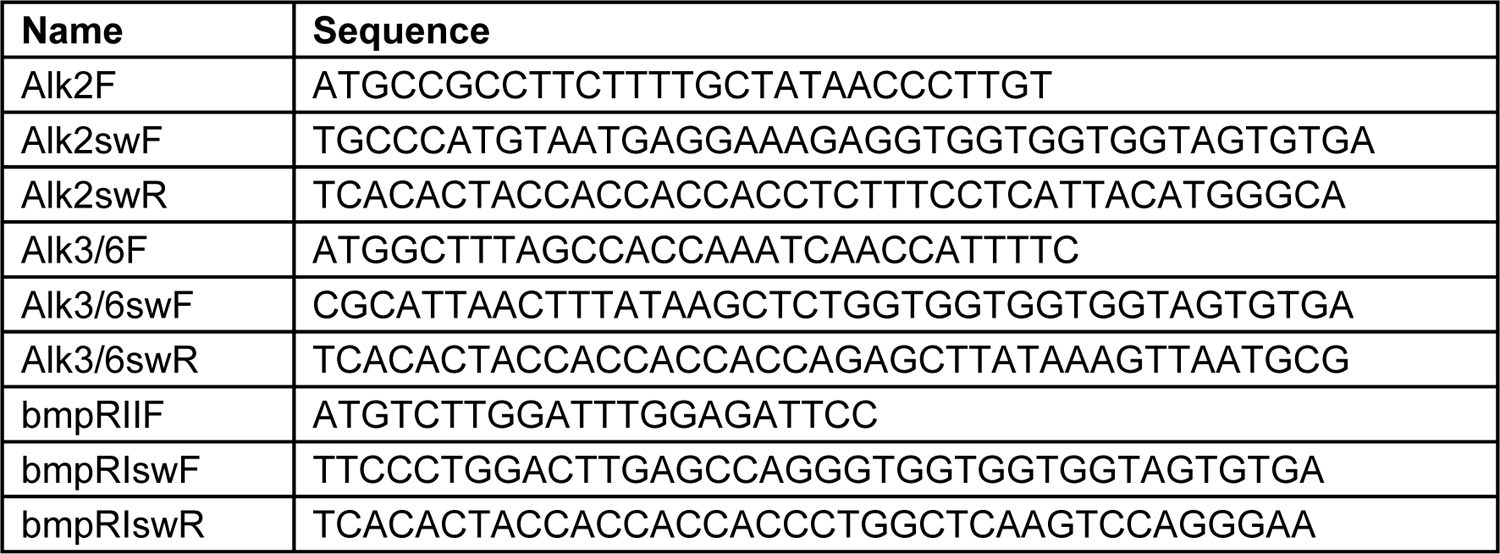
Primers used for cloning dominant-negative BMP receptor fragments

**Supplementary Table 3.**
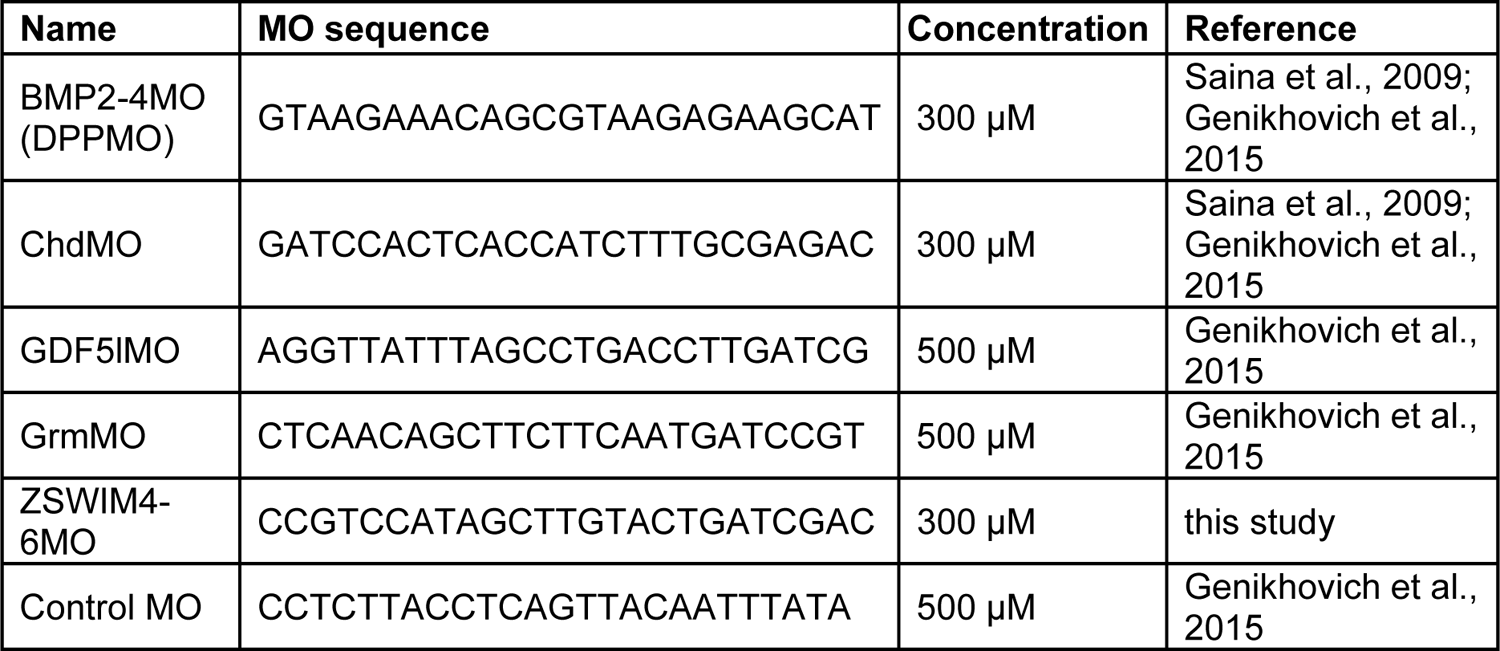
Morpholino sequences

**Supplementary Table 4.**
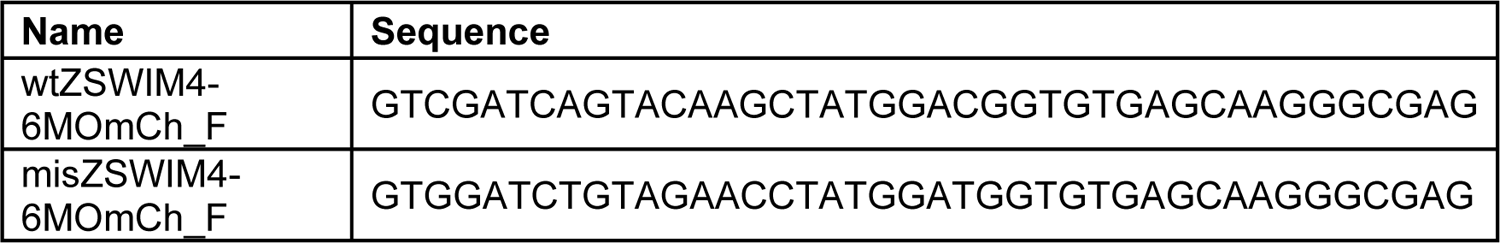
Primer sequences for MO specificity testing

**Supplementary Table 5.**
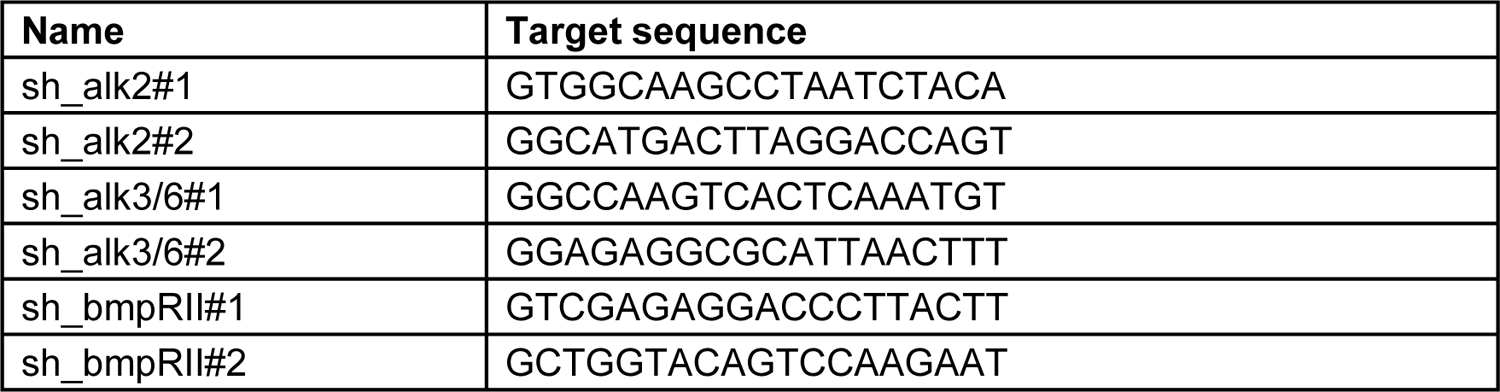
Short hairpin RNA targets

**Supplementary Table 6.**
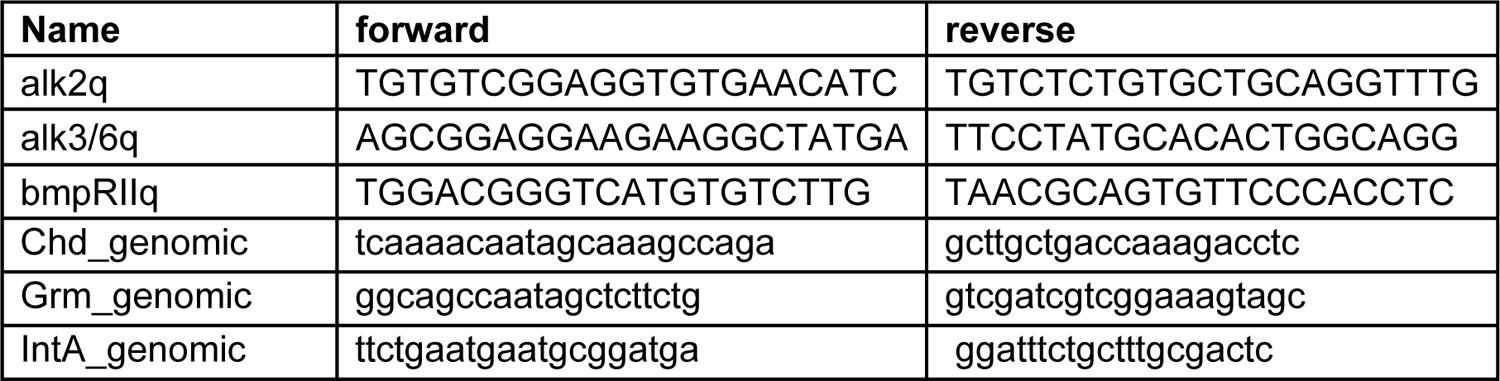
qPCR primers

**Supplementary Table 7.**
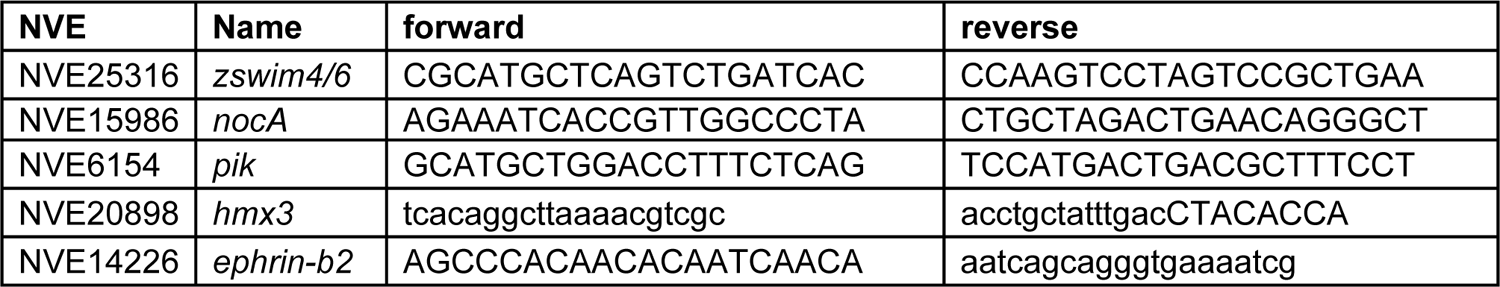
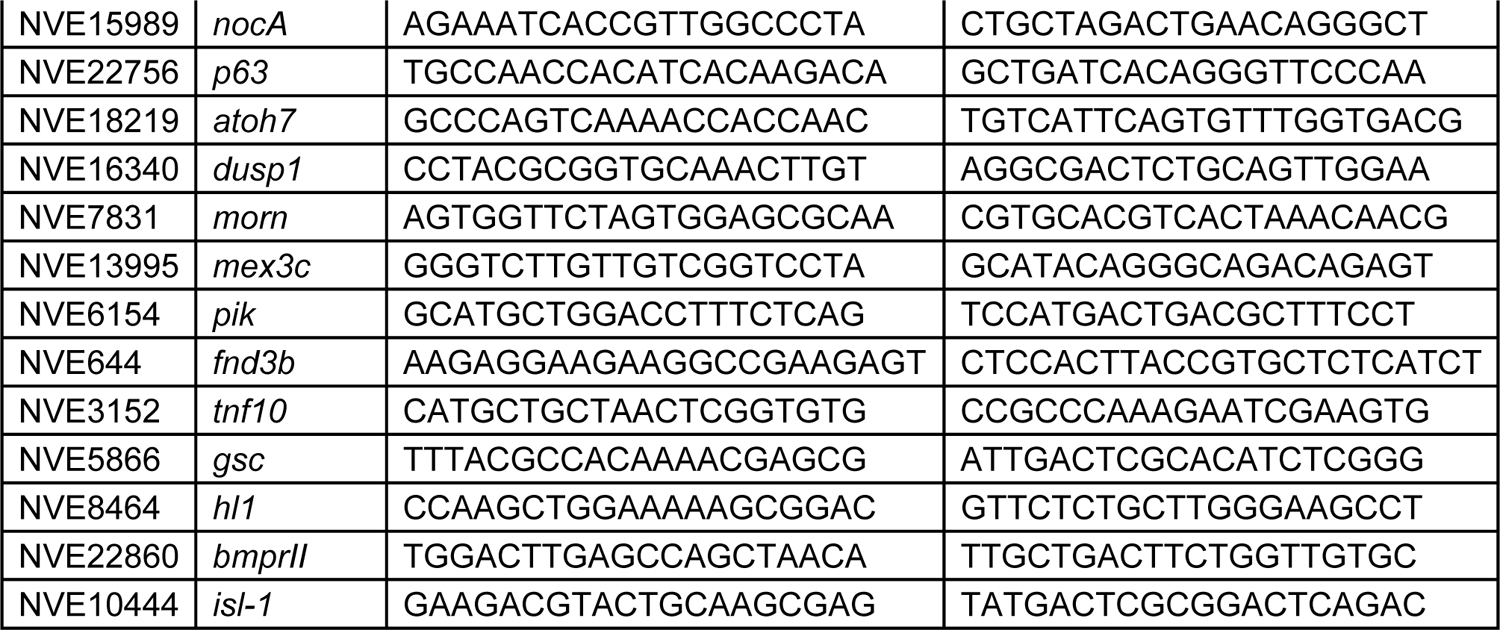
Primer sequences for in situ probes

## Notes

### Competing Interest Statement

The authors have declared no competing interest.

### Summary of Updates

We found a mistake in the legend to Fig. 6 E-F'. In the first version, the legend says "The measurements from Control MO embryos (n=32) and ZSWIM4-6MO embryos (n=33), as well as egfp mRNA embryos (n=24) and zswim4-6 mRNA embryos (n=26)". The n numbers in brackets are, however, incorrect and should be as follows: Control MO embryos (n=10) ZSWIM4-6MO embryos (n=10) egfp mRNA embryos (n=24) zswim4-6 mRNA embryos (n=8)

